# Morphological Integration and Modularity in the Hyperkinetic Feeding System of Aquatic-foraging Snakes

**DOI:** 10.1101/2020.09.16.300400

**Authors:** Daniel Rhoda, P. David Polly, Christopher Raxworthy, Marion Segall

## Abstract

The kinetic skull is a key innovation that allowed snakes to capture, manipulate, and swallow prey exclusively using their heads using the coordinated movement of 8 bones. Despite these unique feeding behaviors, patterns of evolutionary integration and modularity within the feeding bones of snakes in a phylogenetic framework have yet to be addressed. Here, we use a dataset of 60 *µ*CT scanned skulls and high-density geometric morphometric methods to address the origin and patterns of variation and integration in the feeding bones of aquatic-foraging snakes. By comparing alternate superimposition protocols allowing us to analyze the entire kinetic feeding system simultaneously, we find that the feeding bones are highly integrated, driven predominantly by functional selective pressures. The most supported pattern of modularity contains four modules each associated with distinct functional roles: the mandible, the palatopterygoid arch, the maxilla, and the suspensorium. Further, the morphological disparity of each bone is not linked to its magnitude of integration, indicating that integration within the feeding system does not constrain morphological evolution and that adequate biomechanical solutions to a wide range of feeding ecologies and behaviors is readily evolvable within the constraint due to integration in the snake feeding system.

## Introduction

Morphological integration and modularity, defined as the covariances between anatomical traits (integration) and their partitioning into semi-autonomously varying modules (modularity), are key concepts in evolutionary biology and are present at some level in all organisms (Olson & Miller 1958; Wagner 2007; Klingenberg 2008). Morphological integration may be advantageous for maintaining functional associations between traits; however, integration is expected to constrain morphological evolution when the direction of selection is not parallel to the line of least evolutionary resistance, defined by the phenotypic covariance matrix (Wagner & Altenberg 1996; Goswami et al. 2014; Felice et al. 2018; Melo et al. 2016). An integrated phenotype may therefore divert a lineage from evolving across an adaptive landscape along a direct path towards an adaptive peak, consequently deflecting evolutionary responses towards less favorable but more probable regions of morphospace. Modularity thus represents a compromise between complete independence between traits, which promotes evolvability but does not maintain functional associations, and complete integration, which constrains morphological evolution in non-viable ways (Wagner & Altenberg 1996; Goswami 2006). Accordingly, comprehension of the morphological evolution of a clade across macroevolutionary timescales necessarily involves a detailed understanding of the clade’s patterns of integration, as well as the processes that generate them. In this paper, we examine patterns of morphological integration and modularity in the highly kinetic feeding system of aquatic-foraging snake skulls (Fig. 1). The acquisition of a higher degree of kinesis in the elements of the face and mandible compared to other vertebrates is a key evolutionary innovation (Caldwell, 2019) which allowed snakes to radiate into over 3789 living species (Aug 2019, Uetz et al. 2019) encompassing a broad ecological, dietary, and geographic diversity.

**Figure 1.**
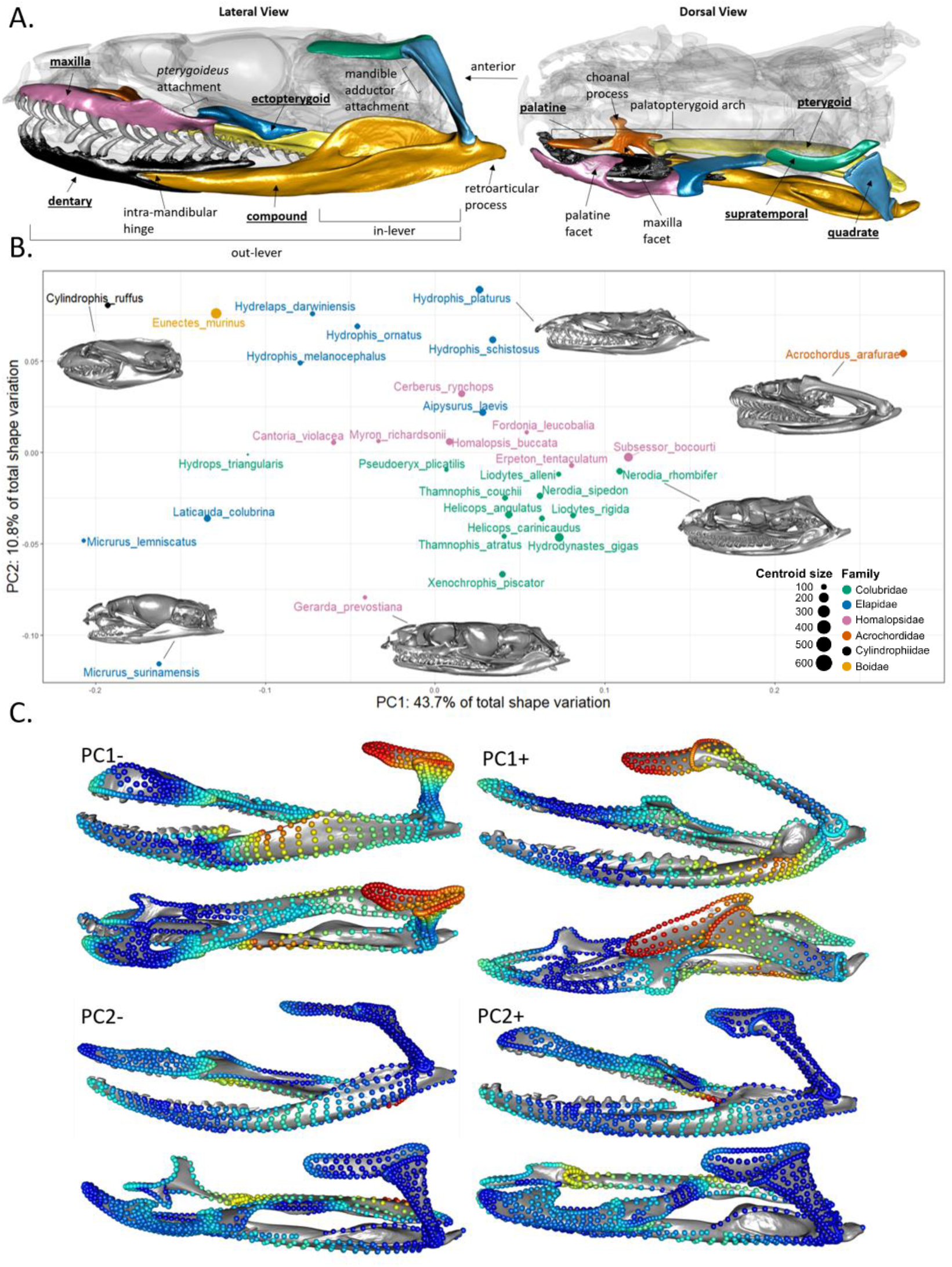
(A) The feeding system of snakes (colored) with the anatomical features referenced in this paper labelled on a specimen of Myron richardsonii (AMNH R111792). The labels of bones studied here are underlined. (B) Scatterplot of the first principal components of a common superimposition of the whole feeding system. Each species is represented as an individual point, the size and color of which corresponds to the centroid size (mm) and taxonomic family (see caption). (C) Shape variation along the first two principal components in lateral (top) and dorsal view (bottom), with landmark colors corresponding to the relative amount that each landmark varies along each PC axis (from PC-to PC+) with red – most variation, and blue – least variation.

The degree and organization of integration among anatomical traits is primarily caused by shared developmental origin and genetic linkages or functional relationships between traits (Olson and Miller 1958; Hallgrimsson et al. 2009). A morphological structure with multiple functions and developmental origins may be both functionally and developmentally modular, not necessarily in overlapping ways (Atchley & Hall 1991; Raff 1996). Disentangling the relative contributions of functional versus development modules will provide a fuller understanding of the macroevolutionary implications of morphological integration because their evolutionary lability reflects the lability and strength of the constraints that populations face in their evolution towards adaptive peaks. For example, if the pattern of evolutionary modularity is caused by a highly conserved suite of developmental pathways, then the macroevolutionary consequences of integration and modularity may be greater than if the same patterns were caused only by functional associations maintained by performance-based selection (Raff 1996). The feeding system of snakes (Fig. 1A) presents an opportunity to study how competing factors translate into patterns of morphological integration and modularity because it is both developmentally and functionally modular in patterns that do not match. If patterns of functional modularity match patterns of evolutionary modularity, then the functional relationships between traits is the primary driver of morphological integration.

Snakes are limbless tetrapods that forage almost exclusively using their heads (Cundall and Greene 2000; Moon et al. 2019). The hyperkinetic skulls of snakes are composed of over 20 bones articulated but unfused with one another, 8 of which are directly involved in feeding (Fig. 1A). The extraordinary kinesis of the feeding system facilitates the independent movements of its individual bones which allows exceedingly large gape sizes such that many snakes can ingest prey much larger than they are (Kardong 1977, 1979). The feeding bones are developmentally modular at least insofar that the different bones are ultimately the results of spatially separated developing cellular populations (i.e., the ossification centers of each bone do not meet to fuse together during development, but see Discussion, Raff 1996; Polachowski & Werneburg 2013; Boughner et al. 2007). Alternatively, the movements of these spatially separated bones must act in concert to successfully forage; snakes must capture, manipulate, and ingest prey exclusively using their head and anterior trunk (Cundall and Greene 2000; Moon et al. 2019). The feeding sequence of snakes can be divided into several segments: prey capture, prey manipulation and repositioning, and swallowing which includes the highly conserved ‘pterygoid walk’ where the teeth of the palatine and pterygoid grasp and hold onto prey while the braincase advances over it (Boltt and Ewer 1964; Cundall and Greene 2000; Moon et al. 2019). The coordinated movement of different groups of bones are required to perform these different functions, forming functional modules. Bones within a functional module share selective pressures associated with their function and would be expected to covary over evolutionary time. Therefore, in the feeding system of snakes there exists a tension between the kinesis and developmental disintegration of bones promoting modularity and the functional dependencies between those bones promoting integration.

Snakes have independently invaded aquatic habits multiple times (over 360 species of snakes use aquatic media, Murphy 2012). The head shape of aquatic foraging snakes has functionally converged in response to the physical constraints related to prey capture under water (Fabre et al. 2016; Segall et al. 2016; 2019). Yet, aquatic-foraging snakes show a large amount of morphological variability along with an exceptional ecological diversity in terms of diet, behavior, and habitat-use (Segall et al. 2020), which may be related to the disparate morphology of their feeding bones (Klaczko et al. 2016). The feeding sequence is highly constrained under water, from prey detection, to the hydrodynamic constraints generated by an accelerated strike (Segall 2019; Segall et al. 2020), to the subjugation and manipulation of slippery (e.g. fish, tapdoles), hard (e.g. crustaceans), and elongated preys (e.g. eels), to the lack of constriction in most species, to swallowing a (sometimes living) neutrally-buoyant prey item (Moon et al. 2019). Piscivorous snakes present some specific morphological features in response to these functional constraints such as numerous, longer, sharper teeth (Savitzky 1983). Yet, the morphological and functional relationships between the different bones involved in the feeding sequence remains poorly studied.

In the present work, we use high-density 3D geometric morphometrics on the 8 bones that compose the feeding system snakes to explore phylogenetically informed patterns of morphological integration and modularity within a large sample of species that share a functional constraint: aquatic foraging. We investigate morphological integration at all levels: within bones, between bones, and within the whole feeding system. To study morphological integration considering the whole trophic system, we compare two superimposition procedures allowing us to analyze separate mobile (articulating) bones at once, and then directly compare *a priori* hypotheses of modularity using the Covariance Ratio effect size (Adams & Collyer 2019). Focusing on a complex system involving the coordination of several morphologically disparate, developmentally disintegrated bones to fulfil a highly constrained behavior (aquatic foraging), this study aims to understand how functional and developmental modularity is translated into evolutionary integration and modularity, and how these patterns affect morphological disparity over macroevolutionary timescales.

## Material and Methods

### Sampling and scanning

Our taxonomic sampling consisted of 60 adult specimens representing 32 species of aquatic-foraging snakes from a wide taxonomic range (Fig. 1, Supplementary Table 1). Specimens came from multiple Museum collections (AMNH, CAS, FMNH) and were carefully chosen to have the mouth closed with no visible deformation or damage. We performed computed microtomography (CT) scans at a resolution between 15-50µm, with the Phoenix v|tome|μCT scanner (General Electric Company, Fairfield, CT, USA) at the AMNH Microscopy and Imaging Facility using a voltage between 100-150kV and current between 130-160mA for a voxel size between 15.6-57.4μm. The 3D reconstruction of the whole skull was performed using the software Phoenix datos|x2 and the subsequent segmentation was done using VGStudioMax v. 3.0 (Volume Graphics GmbH, Heidelberg, Germany). The dentary, compound (here defined as the portion of the mandible posterior to the intramandibular hinge, as in Anjelkovic et al. 2016, 2017), quadrate, supratemporal, pterygoid, ectopterygoid, palatine, and maxilla from the left side of each specimen were digitally separated from the whole skull in GeomagicStudio (3D Systems, Rock Hill). To facilitate the deployment of surface sliding semilandmarks, each bone was cleaned in Geomagic so that small holes were covered, teeth were removed, and surfaces were smoothed following the procedures suggested by Bardua and colleagues (2019a).

### 3D Geometric Morphometrics

We used a high-density 3D geometric morphometric approach (1335 total landmarks across all bones) using both anatomical and semilandmarks, to quantify the shapes of each bone (Adams et al. 2004, 2013; Dumont et al. 2016; Goswami et al. 2019). Anatomical landmarks and curve semilandmarks were placed on each bone using IDAV Landmark Editor and MorphoDig (Supplementary Fig. 1; Wiley et al. 2005; Lebrun & Orliac 2017). Using the function ‘placePatch’ in the R package *Morpho*, surface landmarks were projected onto each separate bone from a template specimen following the precautions suggested by Bardua and colleagues (2019a; Schlager 2017). Disparity in the shapes of the choanal and maxilla facet of the palatine obstructed the projection of surface landmarks, so the shape of the palatine was represented only with anatomical and curve landmarks. The choanal and maxilla processes were still captured with anatomical and curve landmarks. The curve and surface semilandmarks were allowed to slide to minimize bending energy (Gunz et al. 2005; Gunz & Mitteroecker 2013). We used generalized least-squares Procrustes superimposition with the ‘gpagen’ function in *geomorph* (Rohlf & Slice 1990; Adams and Otárola-Castillo 2013) to analyze shape variation in each bone, individually.

### Shape variability, phylogenetic signal, and allometry

To define lines of least resistance, we used principal component analyses (PCA, ‘gm.prcomp’ function in *geomorph*) to extract and visualize the main axes of variation for each bone and the whole feeding system (Adams and Otárola-Castillo 2013; see below for an account of how bones were combined to assess the feeding system as a whole). Thin-plate spline deformations applied on meshes were used to visualize the shape variation associated with each axis (‘tps3d’ function in *Morpho*). Using the phylogeny of Pyron and Burbrink (2014) pruned to our dataset (Supplementary Table 1), we tested for a phylogenetic signal in each bone (‘physignal’ function in *geomorph*) to assess whether the phylogenetic relationships between species was related to their morphology (Adams 2014; Adams and Otárola-Castillo 2013). A significant phylogenetic signal was found in each bone (P<0.01, K<0.59, Table 1), so all subsequent analyses were performed in a phylogenetically informed context. To test for the effects of evolutionary allometry on shape, we performed phylogenetic generalized least squares (PGLS) analyses on the Procrustes coordinates and the log-transformed centroid size as a covariate using the ‘procD.pgls’ function in *geomorph*.

### Modularity analyses and superimposition protocols

As employed here, both eigenvalue dispersion and two-block partial least squares measure covariance in shape within and between bones independent of their relative sizes and positions in the mouth (discussed below). However, in the case of the feeding bones of snakes, the relative positions, orientations, and sizes of bones are immensely important to the functional relationships between bones and therefore morphological integration and modularity. For example, a larger gape in many taxa is accomplished by the backwards rotation of the quadrate, such that the quadrate points posteriorly (see *Acrochordus arafurae*, Fig. 1B) rather than orthogonally to the mandible (as in *Cylindrophis ruffus*, Fig. 1B). Further, patterns of ontogenetic allometry in some macrostomatan snakes involve backwards rotation of the quadrate and positive allometry of the jaws, supratemporal length, and quadrate length (Scanferla 2016; Palci et al. 2016), facilitating ontogenetic niche shifts in some species (Vincent et al. 2007; Mushinsky 1982). This allometric axis of shape variation, affecting separate component parts (i.e., potential modules) of the snake feeding system simultaneously, has been demonstrated to be functionally consequential, evolutionary labile, and adaptive (Esquerré et al. 2017, and Sherratt et al. 2019 report heterochronic shifts facilitating dietary shifts), suggesting that it may influence evolutionary integration within the snake skull. A shared coordinate system that maintains the relative sizes and positions of the bones is therefore desirable to understand the patterns of integration and modularity. We employed two superimposition strategies to achieve this goal: a common superimposition and ‘matched’ local superimpositions.

The common superimposition consisted of performing a GPA on all of the bones at once in their original CT scanned positions, treating them as if they were a single structure (GPA_all_) because in that position the bones all retain their relative sizes and are in a natural resting position relative to one another that is largely repeatable (it is worth noting that no ‘true’ anatomical position exists in a kinetic system, Collyer et al. 2020). Mobility between the bones in a living snake is comparatively small relative to their overall position in this resting orientation (Watanabe et al. 2019, Supplementary Fig. 2). Other authors have adopted a similar strategy for analyzing the shape of the entire feeding apparatus (e.g., Watanabe et al. 2019; Palci et al. 2016; Klaczko et al. 2016; Souto et al. 2019; Silva et al. 2018; Murta-Fonseca et al. 2019; dos Santos et al. 2017) and additional studies have superimposed non-rigid structures together in other organisms when taking appropriate precautions (Adams 1999; Rohlf & Corti 2000; Adams & Rohlf 2000; Adams 2004). Each specimen was μCT scanned with the mouths completely closed; only specimens in neutral poses were included, where articulating bones were positioned directly adjacent to each other. Even though none of the individual bones are fused with one another, numerous soft tissue connections reduce rotational degrees of freedom of each bone. We also corrected for intraspecific variation by averaging landmark configurations per species, which further minimizes the variation due to mobility alone. The position and respective centroid size of each bone in the mean shape (of all species) resulting from this procedure was computed and used in the local superimposition procedure.

Because the relative positions of the elements could still vary non-repeatably because of the death position in which the specimen was preserved, we also adopted a second: matched local superimposition. This local superimposition procedure first consisted of performing a GPA for each bone separately. The superimposed landmark coordinates of each bone were then translated, rotated, and scaled to fit its corresponding bone of the mean shape from GPA_all_ (Fig. 2). This procedure allowed us to have a more homologous positioning and respective size for all species and to preserve the intrinsic, pure shape variation of each individual bone. Further, the mean landmark configuration of the common superimposition, for which the local superimpositions were matched onto, was biologically realistic; since each specimen included was in a neutral pose, the landmark configurations average to a shape reflects a specimen in a neutral pose (Fig. 2), and specimens are scattered around the origin of the PC morphospace suggesting that the mean configuration is natural and plausible (Fig. 1B). The interspecific positional and rotational variation between species may be large enough that it confounds interspecific patterns of pure shape variation within bones. The functional and developmental processes that govern the positional variation in bones may not be the same exact processes that govern shape variation in individual bones, and therefore support for alternative patterns of modularity may disproportionately reflect processes that control the positional variation in bones when only considering a common superimposition. The local superimposition procedure set landmark coordinate configurations in a coordinate space that was biologically realistic and reduced the positional and rotational variation due to mobility so that we could analyze patterns of modularity in the whole system while only considering pure shape (co)variation within the feeding system.

**Figure 2.**
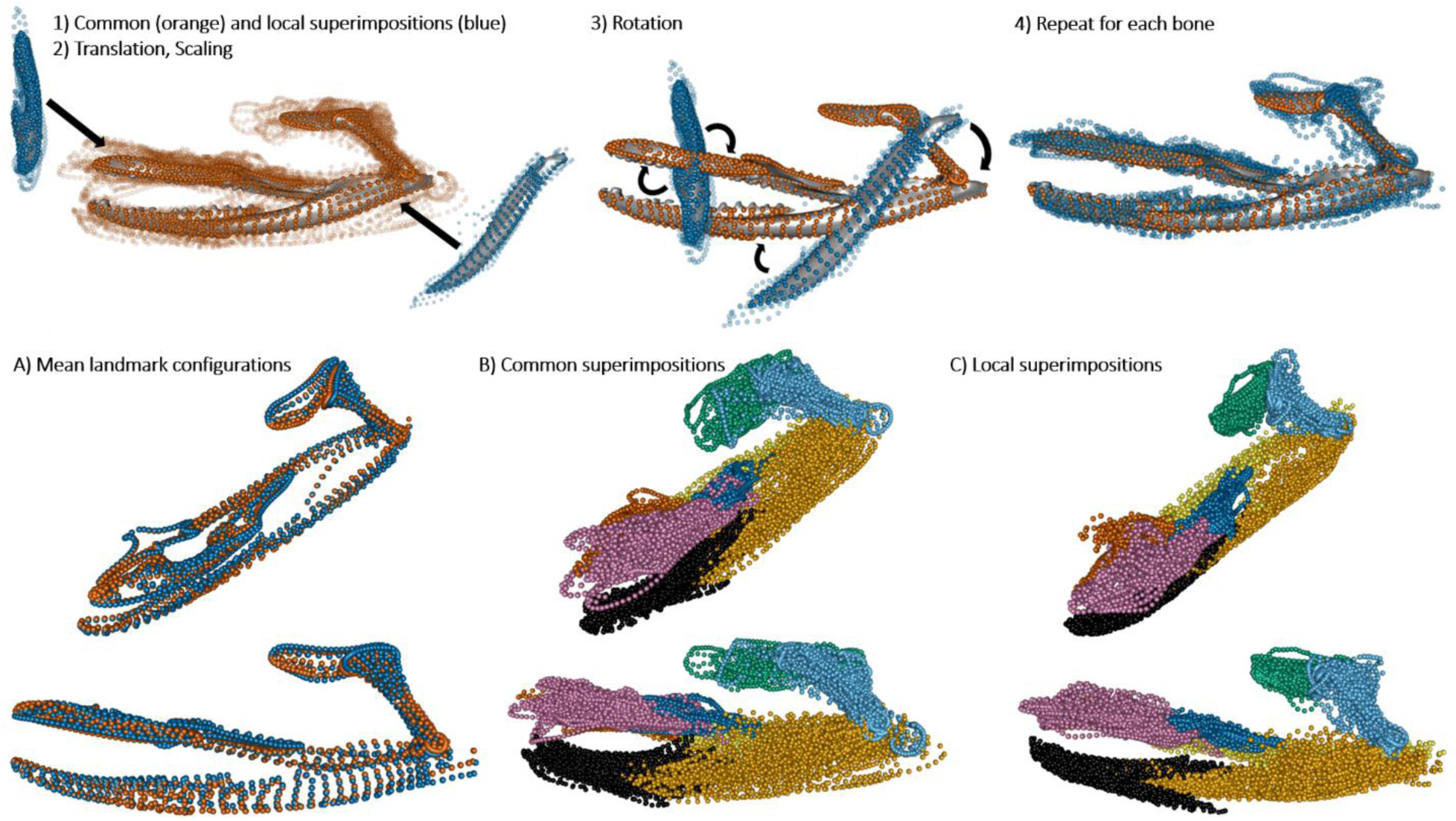
Workflow of the local superimposition procedure: 1) Procrustes superimpositions of the whole feeding system (i.e. common superimposition; GPA_all_, orange), and of each individual bone (i.e. local superimpositions; blue) are performed. The transparent landmarks are individual species after their respective superimpositions. 2) Each local superimposition dataset is translated and scaled to the mean centroid size and position of its corresponding bone in the global mean shape from GPA_all_ and (3) rotated. 4) Final dataset for the local superimposed procedure. (Below) A) Superimposition of the mean configurations of the common (orange) and local superimpositions dataset (blue). Complete landmark datasets for the common (B) and local (C) superimpositions procedures (color code matches Fig. 1).

The two superimposition procedures are complementary: the local superimpositions account for pure shape variation in each bone and the common superimposition emphasizes (co)variation related to the overall configuration of the feeding apparatus. By comparing results from the two procedures, we bracket the ‘true’ pattern of morphological, evolutionary modularity. If both superimposition procedures support similar hypotheses of modularity, then it is unlikely that the pattern arises because of arbitrary differences due to positioning of the mobile elements.

### Morphological integration and modularity analyses

We used three methods to analyze the pattern of morphological integration and modularity. Relative eigenvalue standard deviation (eigenvalue dispersion) was employed to measure the overall degree of morphological integration within each bone and then the feeding system as a whole (Pavlicev et al. 2009). Phylogenetic two-block partial least squares (2BPLS, ‘phylo.integration’ function in *geomorph*) analyses were used to assess morphological integration between each pair of bones (Rohlf & Corti 2000; Adams & Felice 2014; Adams & Collyer 2016). The Covariance Ratio was used to test *a priori* hypotheses of modularity based on the whole feeding system (Adams 2016, Adams & Collyer 2019).

Eigenvalue dispersion was calculated from a singular value decomposition of the covariance matrix of the Procrustes-superimposed landmark configurations for each bone. Higher eigenvalue dispersion values correspond to larger degrees of morphological integration because a smaller number of eigenvectors will explain a larger portion of total correlated shape variation in more-integrated structures (Pavlicev et al. 2009; Goswami & Polly 2010a). Eigenvalue dispersion values range between 0 and 1 and are comparable across datasets (e.g., different bones, Pavlicev et al. 2009).

The degree and significance of morphological integration between each pair of bones was quantified using phylogenetic 2BPLS and its effect size (Rohlf & Corti 2000; Adams & Felice 2014; Adams & Collyer 2016). For each pair of significantly integrated pairs of bones (p-value < 0.05) we describe shape variation along the primary axis of covariation (PLS1) to determine which anatomical structures contribute to covariation (Fig. 4). To test if the magnitude of integration constrains morphological diversity, we conducted two least-squares linear regression analyses with Procrustes variance values (i.e., morphological disparity) against 1) eigenvalue dispersion values (i.e., within-bone integration) and 2) the average 2BPLS effect size for each bone (i.e., between-bone integration) (Fig. 3).

**Figure 3.**
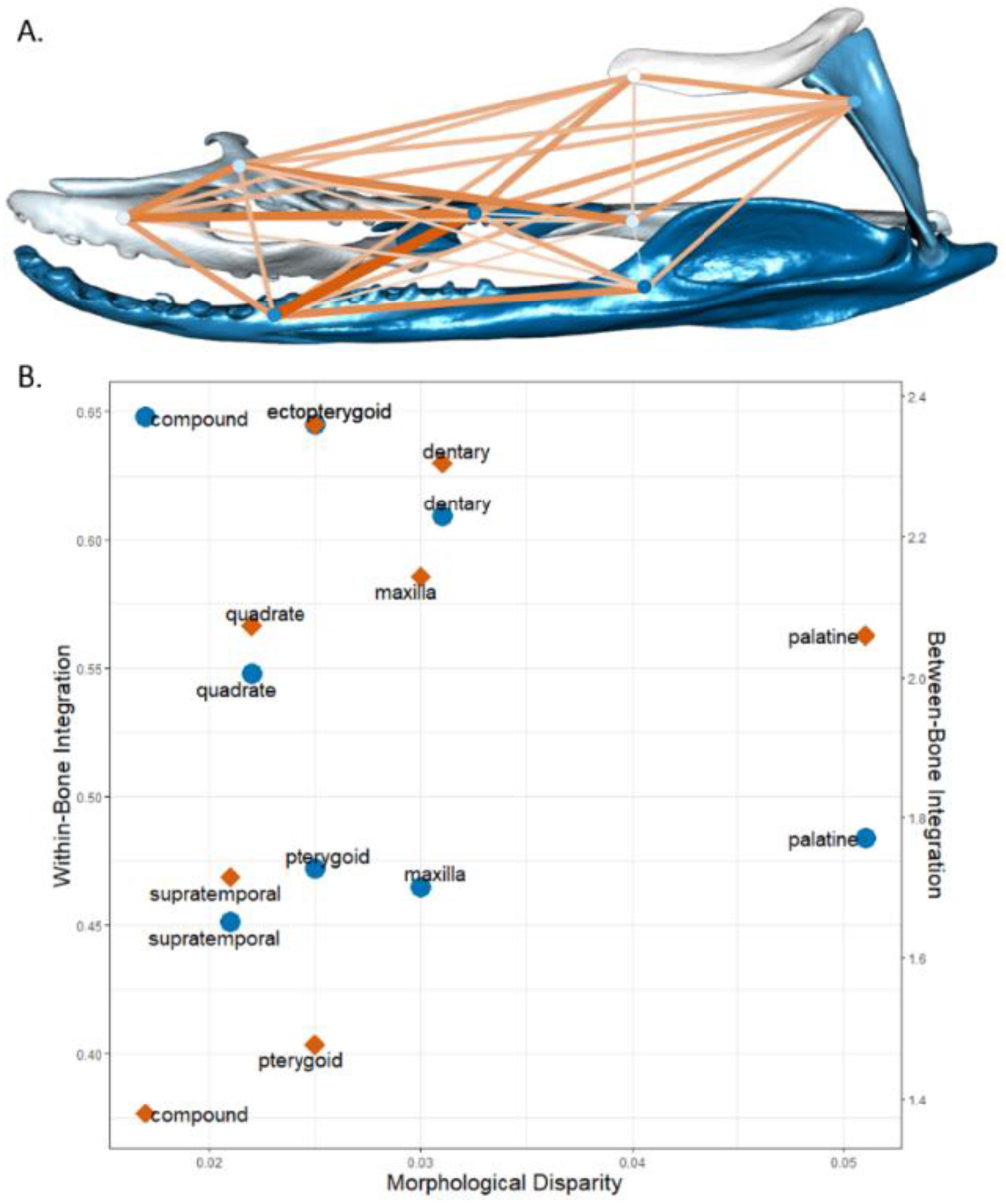
A) Network graph of within and between-bone integration in the feeding apparatus. Within-bone integration (eigenvalue dispersion values) is represented by the bone color: the darker, the stronger. Covariation between bones (2BPLS effect size; z-PLS) is represented by the color and width of the line connecting each pair of bones: wider and darker connections correspond to higher effect sizes and stronger morphological integration between the bones (values in Table 1, Supplementary Tables 3-4). B) Scatter plots of morphological disparity (x-axis) with within-bone integration (eigenvalue dispersion, blue points, left y-axis) and between-bone integration (average all the z-PLS for each bone, orange points, right y-axis). Both regression analyses were not significant.

**Figure 4.**
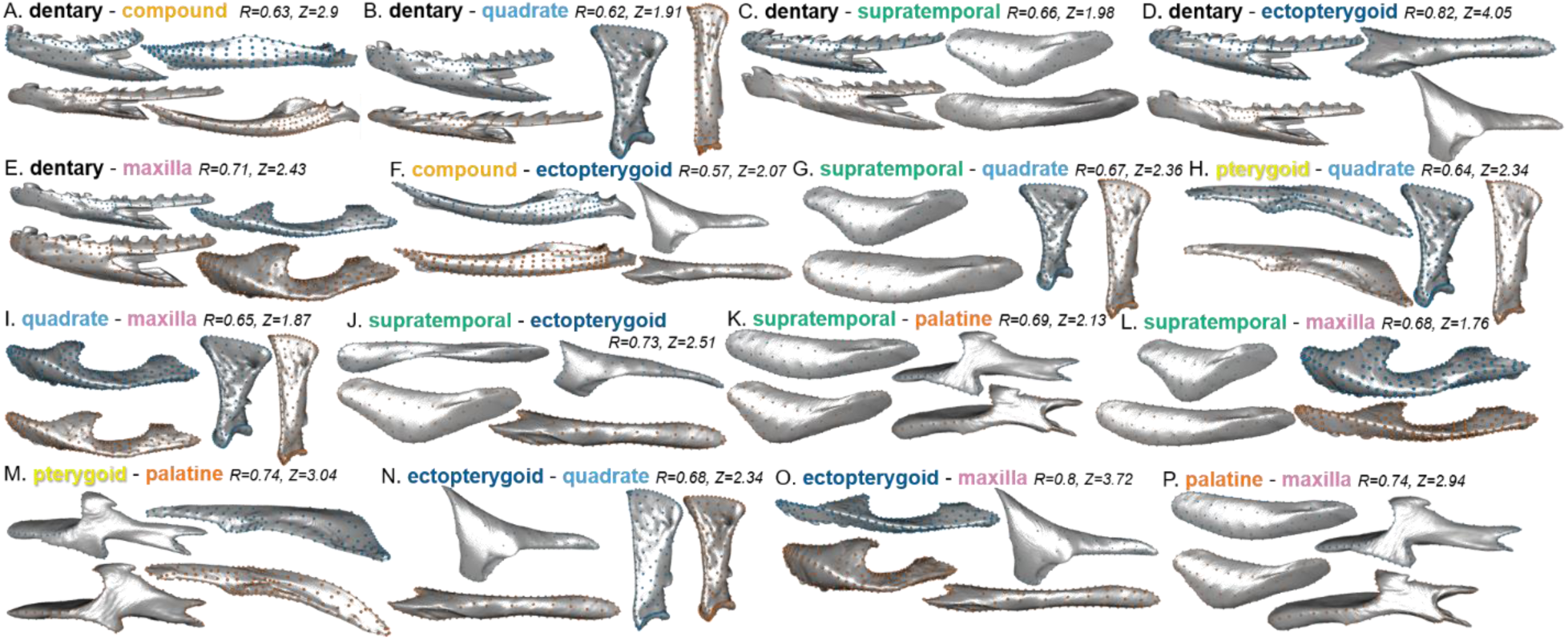
Figure 4. Pairwise shape covariation in snake feeding bones along PLS1 from each significant phylogenetic 2B-PLS. (blue: PLS1-; orange: PLS1+)

For both superimposition methods, 21 *a priori* hypotheses of modularity were tested using the CR (Adams 2016; Adams & Collyer 2019). Each hypothesis of modularity was based on combinations of associations of bones that would be expected to covary in certain functional or developmental contexts (Supplementary Table 2).

To visualize the major axes of correlated shape variation within each module of the most supported hypothesis, we performed phylogenetic PCAs (pPCA, Revell 2009, Adams & Collyer 2018) on each module (‘gm.prcomp’ in *geomorph* with the ‘Transform’ and ‘GLS’ parameters set to ‘True’). Bones within each module were superimposed together. These per-module superimpositions weren’t used for any statistical analysis and were only used for visualization purposes. Shape variation along pPC1 was visualized because it is the axis of most correlated shape variation within each module while accounting for phylogenetic non-independence.

## Results

### Shape variation and covariation of individual bones

Significant allometries were found in the quadrate, supratemporal, palatine, and maxilla but explained only a small portion of the shape variation (R^2^=0.6-0.8) except for the quadrate (R^2^=0.26, Supplementary Table 3). There was no significant allometry in the dentary, compound, pterygoid and ectopterygoid. The largest eigenvalue dispersions (i.e. within bone integration) were reported in the compound (0.648) and ectopterygoid (0.645) and the smallest in the supratemporal (0.451) and maxilla (0.465) (Supplementary Table 3, Fig. 3). Linear regression analysis showed that the degree of morphological integration within each bone did not significantly relate to morphological disparity (p-value=0.44, Fig. 3). Our 2BPLS analyses recovered significant integration between 16 out of 28 possible pairs of bones (Supplementary Table 4, 5, Fig. 4). The pterygoid is only significantly integrated with the quadrate and palatine, and the compound is only significantly integrated with the dentary and ectopterygoid. The palatine is also integrated with less bones (3) than any of the other bones (each is integrated with 5 other bones) (Fig. 4, Supplementary Table 3, 4). The more strongly integrated pairs are ectopterygoid/dentary (r-PLS=0.822, Fig. 4D) and the ectopterygoid/maxilla (r-PLS=0.806, Fig. 4O). Descriptions of shape variation along PLS1 are provided in the Discussion and shown in Fig. 3. The average 2BPLS effect size for each bone did not significantly relate to morphological disparity (linear regression analysis, p-value=0.173, Fig. 3b).

### Shape variation of the whole feeding apparatus

Significant allometry was recovered for both the common and local superimposition datasets containing all bones (p-value=0.001, R^2^=0.13 and p-value=0.018, R^2^=0.07, respectively). The common superimposition showed a larger eigenvalue dispersion value (0.447) than the local superimposition dataset (0.352).

The first principal component (PC1) of the common superimposition dataset (Fig. 1B, C) accounts for 43.7% of the overall shape variation in the feeding apparatus of snakes and is mainly driven by the orientation and length of the quadrate, as well as slenderness of the mandible and maxilla. Shapes at the positive extreme of PC1 (PC1+) had a longer and backwards-rotated quadrate, and elongated, slender mandibles and maxillas. A ventrally bowed mandible and prominent palatine processes were characteristic of shapes at PC2-. Colubridae grouped together except *Hydrops triangularis* (Colubridae) which was separated along PC1, and Homalopsidae are grouped along PC2, except the crab-tearing specialist *Gerarda prevostiana* (Homalopsidae, Jayne et al. 2002; Jayne et al. 2018). The elapids showed substantial variation along both PC axes and drive the variation of PC2, with the true sea snakes (Hydrophiinae) grouping at PC2+ and the semi-aquatic coral snakes (Micrurinae) at PC2-. *Acrochordus arafurae*’s exceptionally elongated mandible and quadrate drive the variation along PC1+, while species on PC1-have a short and almost horizontally positioned quadrate and a dorsally expended prearticular process. Shape variation along PC2 is smaller, and mainly carried by variation in the palatine-pterygoid joint. Species at PC2-have a dorso-lateral expansion of the posterior part of the pterygoid and a simple shape of the palate-pterygoid articulation while species at PC2+ have a slender pterygoid with a complex articulation shape.

### Modularity in the whole feeding apparatus

The most supported hypothesis (i.e., the most negative Z_CR_ value, meaning the strongest modular signal) for both common and local superimpositions datasets described two modules (Fig. 5). The most supported hypothesis from the common superimposition (H1) described dorso-ventral modularity with mandibular (dentary, compound) and non-mandibular elements as separate evolutionary modules. The most supported hypothesis from the local superimposition dataset (H4) was also composed of two modules, one with the dentary, compound, maxilla and quadrate and the other with the supratemporal, ectopterygoid, pterygoid, palatine, describing a latero-medial pattern modularity. Hypothesis 15 (H15, Fig. 6, Supplementary Table 2, 5) was the third and second most supported hypothesis in the common and local superimposition dataset respectively (Fig. 5). H15 described 4 modules: the mandible, a module with the pterygoid, palatine, and ectopterygoid, the quadrate and supratemporal as a module, and the maxilla as an independent module. Complex hypotheses of modularity (i.e., complete modularity, H21) were not as strongly supported as hypotheses with four or less modules (Supplementary Table 5).

**Figure 5.**
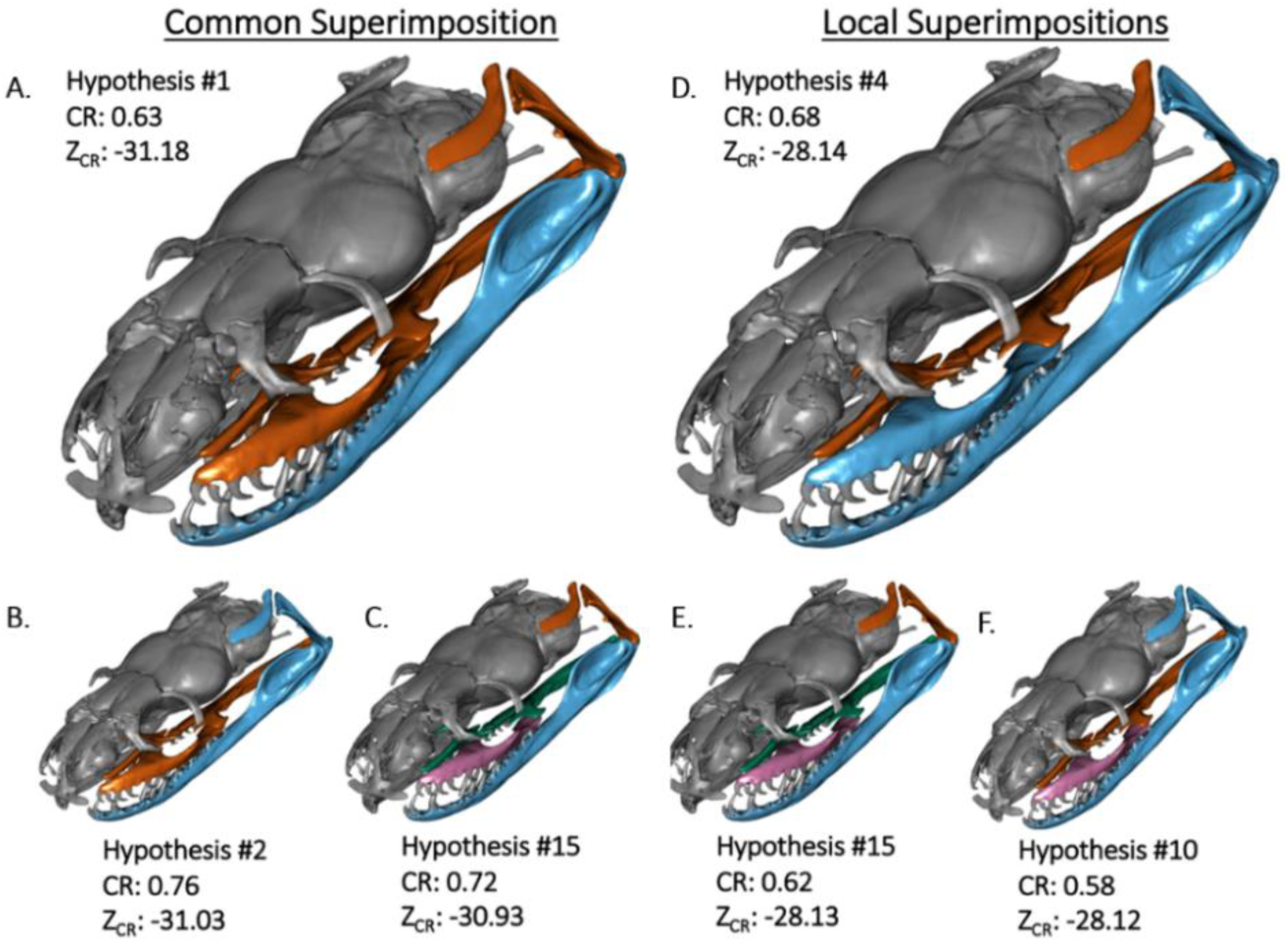
The three most supported hypotheses of snake feeding bone modularity for each superimposition method. A, D – most supported hypothesis, B, E – second most supported, C, F – third most supported. Each group of similarly colored bones describes a module. The lower the effect size (Z_CR_), the stronger the modular signal, and the more supported the alternative hypothesis. The order of most support is supported by pairwise effect size p-values (Supplementary Fig. 11, 12). Note that hypothesis #15 (with four modules) is the only hypothesis supported strongly by both superimposition methods.

**Figure 6.**
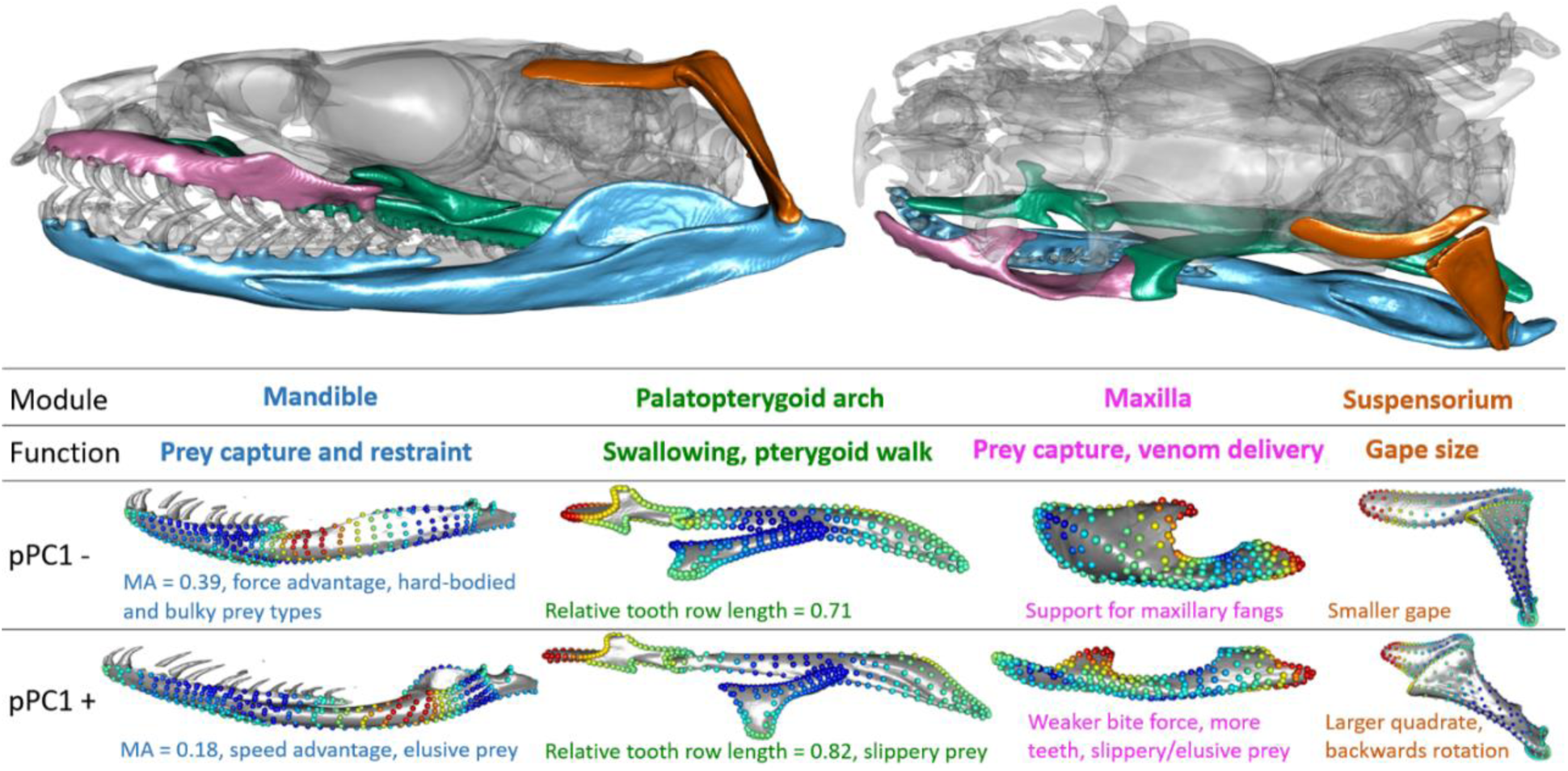
A four-module hypothesis (H15) where each module is associated with a distinct functional selective pressure is strongly supported by both superimposition procedures. (Above) Snake skull with the different modules highlighted in different colors in lateral (left) and dorsal (right) view. (Bottom) Per-module shape variation and their functional consequences along phylogenetic PC1. Relative tooth row length was calculated as (length of palatine tooth row + length of pterygoid tooth row) / total anterior-posterior length of the palatopterygoid arch. The color of landmarks represents the magnitude of its variation along pPC1 with red dots varying the most.

## Discussion

In the present work, we quantified and compared morphological variability of eight bones that work jointly to fulfill a fitness relevant function (i.e. feeding), at different levels of organization, from covariation within individual bones to patterns of integration and modularity considering all bones, in a phylogenetically informed context. Our results suggest that the feeding apparatus of snakes is highly integrated, predominantly driven by functional relationships between the bones. Our most supported hypotheses describe the mandible and the palatopterygoid arch (i.e. palatine, pterygoid) as two separate modules, with the maxilla and quadrate either integrated with the cranial elements (common superimposition) or the mandibular elements (local superimpositions). The hypothesis with the strongest support from both datasets, H15, contains an integrated palatopterygoid arch (including the ectopterygoid), the maxilla as an independent module, the quadrate and supratemporal as a module, and the mandible as a module. This considerable degree of integration is interesting considering the extreme kinesis of the feeding system, a factor we would generally expect to promote modularity because the bones are not physically coupled with one another and have the freedom to move (and evolve) independently. The high degree of integration within the feeding system supports functional relationships between bones as the primary driver of integration, as the synchronized movements of different combinations of trophic bones is necessary for successful foraging (further discussed below, Cundall and Greene 2000; Moon et al. 2019). These findings also corroborate a recent study by Watanabe and colleagues (2019) who found that the crania of snakes are highly modular, except for the palatopterygoid arch. Although only four bones overlap between their study and ours (the distal quadrate ‘jaw joint’, maxilla, pterygoid, and palatine), we report generally consistent patterns of modularity in the most supported hypotheses (Fig. 5, 6), with an integrated palatopterygoid arch separated from the maxilla and quadrate. Although the maxilla and quadrate are a part of the same module in our most supported hypotheses, they show a moderately low degree of integration, and are a part of separate modules in many other highly supported hypotheses, as is discussed below. Moreover, Watanabe and colleagues (2019) speculated that cranial kinesis may promote integration; we find exceptionally strong integration when considering the most mobile elements of the hyperkinetic snake skull.

### Functional and morphological integration in the feeding system

Our most supported hypotheses of modularity are consistent with a pattern that arises from functional aspects of feeding. Prey ingestion is accomplished in most advanced snakes via the ‘pterygoid walk’, involving the coordinated movement of the palatine and pterygoid bones (palatopterygoid arch) (Boltt and Ewer 1964; Cundall 1983), accordingly, there are shared functional selective pressures between these bones. During the pterygoid walk, the palatopterygoid arch teeth grasp prey while the braincase advances over it. This function is crucial for foraging in snakes and its performance is especially fitness-related in macrostomatan taxa that ingest large prey, as snakes are vulnerable to predators during the pterygoid walk (Cundall and Greene 2000). In aquatic-foraging snakes, selective pressures associated with swallowing may be exacerbated due to the lubriciousness of prey, or the fact that there may not be a substrate to anchor onto during the pterygoid walk. The palatine and pterygoid were integrated in our 11 most supported hypotheses from the common superimposition, and 8 most supported from the local superimposition dataset. In fact, the hypothesis of complete modularity besides the palatopterygoid arch (H19) was much more supported in both datasets than the hypothesis of complete integration besides the palatopterygoid arch, indicating that the palatopterygoid arch contributes substantially to covariance patterns when considering the whole feeding apparatus. According to 2BPLS analyses neither the palatine nor pterygoid are integrated with more than three different bones. Andjelkovic and colleagues (2017), who also found low integration between the pterygoid and other bones, hypothesized that the pterygoid’s functional optimization prevents it from covarying with other bones. Considering the pterygoid’s integral role in prey ingestion (hence the ‘pterygoid walk’) and its physical entanglement with multiple other bones either through articulations or muscle attachments, functional optimization may explain the pterygoid’s high integration when considering the whole trophic system but not in 2BPLS analyses. The pterygoid and palatine are also the first bones in the feeding system to ossify (Sheverdyukova 2018; Khannoon & Evans 2015; Polachowski & Werneburg 2013; Werneburg et al. 2015), which may promote integration between them (and disintegration between the palatopterygoid arch and the rest of the system) either because of additional, shared opportunity for bone remodeling during development, or a shared influence of morphogens expressed during the palatopterygoid arch’s early ossification that the other bones do not face.

Strong functional selective pressures drive integration within the mandible as well. The mandible, particularly the compound bone, is directly involved in multiple functional modules. The jaw adductor, responsible for closing the lower jaw and necessary for prey capture, originates on the anteroproximal quadrate and inserts on the mandibular fossa of the compound (Johnston 2014). The *pterygoideus*, which moves the palatopterygoid arch ventromedially and the dentary row dorsolaterally during prey ingestion, attaches on the retroarticular compound process and either the ectopterygoid or maxilla (Cundall 1983; Jackson 2003; Johnston 2014, Fig. 1A). Therefore, we would expect the compound to be integrated with the dentary, quadrate, ectopterygoid, and maxilla if functionally relevant muscles solely cause morphological integration. Yet 2BPLS results reveal there is significant integration only with the dentary and ectopterygoid. The PC morphospace and 2BPLS results both suggest that selection for mechanical advantage (MA) is the primary driver of integration within the mandible. The main component of variation in the compound bone (PC1), that accounts for almost 65% of the variation, describes the ratio of the in-lever to the out-lever (Fig. 1A). PC1 of the dentary describes variation in its slenderness and relative length (Supplementary Fig. 3). Shape variation along PLS1 of the dentary and compound mirror this shape variation (Fig. 4a). Taken together, the main axes of variation (PC1) and covariation (PLS1) in the dentary and compound cumulatively describe variation in the MA of the species. Species with a low MA (a longer out-lever compared to in-lever) have a speed advantage and are adapted for capturing elusive prey, and species with a high MA will have a force advantage and are adapted for capturing hard-bodied or bulkier prey (Hampton 2011; Wainwright and Richard 1995; Mori and Vincent 2008). Selection for MA almost certainly promotes integration between the dentary and compound because both separate bones contribute to MA. Further, jaw MA has been related to dietary niche (Mori and Vincent 2008; Hampton 2011), meaning that selection for dietary specializations may promote integration within the mandible. The dentary and compound were significantly integrated (2BPLS) and were a part of the same module in our four most supported hypotheses of modularity (Table 4). This substantial integration within the mandible is functionally relevant, and is particularly interesting considering the evolution of the intra-mandibular hinge that can be considered a form of developmental disintegration in that ossification centers of the dentary and compound do not fuse, a process we would expect to promote modularity, especially if inhibitory signaling obstructs fusion of the ossification centers (Raff 1996). The anatomical liberation between component parts of the mandible is functionally adaptive, as it allows greater mobility and a larger gape (Kardong 1977); in the mandible we find strong morphological integration directly from developmental disintegration.

### Dietary niche and modularity

If the relative importance of different functional modules relates to dietary niche, then it is reasonable to assume that patterns of modularity are influenced by the selective pressures associated with both prey properties and feeding behaviors. For example, eels require different manipulation skills than crabs, however capturing and restraining hard-bodied crabs may require a stronger bite force than required to capture an eel. Consequently, an eel specialist may exhibit a covariance structure such that features associated with prey manipulation are more strongly integrated than features unique to prey capture, and vice versa in a crab-specialist taxon.

The *retractor pterygoideus* which originates on the braincase and inserts on the medial palatine along the choanal process, is responsible for advancing the braincase over the prey during the pterygoid walk. The shape of the choanal process on the palatine (C8, C9, Supplementary Fig. 1), is highly variable along PC2 of the common superimposition, and both PC axes of the palatine morphospace (Supplementary Fig. 9). The venomous, pelagic *Hydrophis platurus* completely lacks a choanal process but contains an elongated retroarticular process, providing substantial area for attachment of the *pterygoideus* muscle. Stimulation of the *pterygoideus* muscle induces outward rotation of the dentary tooth row (Cundall 1983). The primary role of the mandible in many taxa during ingestion is to keep prey pressed onto the teeth of the palatopterygoid arch. In a venomous,, pelagic sea snake adapted for elusive and neutrally buoyant prey, selective pressures for prey ingestion may be stronger on morphological features relevant to prey manipulation and handling rather than advancement of the braincase over bulky, less mobile prey, consequently creating different patterns of integration involving areas of attachment for these or other cephalic muscles. The degree to which prey properties, particularly shape or bulkiness, affects patterns of integration in the feeding system of snakes by reorganizing the relative importance of different, overlapping, functional modules should be investigated in further detail. Additionally, the alternative modes of prey ingestion adapted to a broad range of prey types (e.g., mandibular raking, Deufel & Cundall 2003, or sawing, Kojima et al. 2020, or tearing prey, Jayne et al. 2018) illustrates the interspecific variation in myological and functional relationships between the feeding bones. Future studies should examine intraspecific modularity in a select number of taxa at phylogenetically and ecologically informative positions to control for this variation in functional modularity, and to further incorporate within-bone modularity into hypotheses of modularity of the whole feeding system. Accompanying these morphometric analyses with empirical measurements of functional performance (e.g., manipulation and swallowing durations) will provide a better understanding of the factors shaping patterns of integration and modularity in the feeding bones of snakes, specifically how shifting functional relationships between anatomical regions translate into phenotypic covariance.

### Morphological integration in the maxilla and upper jaw

The placement of the maxilla in either the palatopterygoid arch module, the suspensorium module, or as an independent module is not apparently clear. Cundall (1983) argued for a medial swallowing functional module, composed of the palatopterygoid arch, and a lateral prey capture functional module, composed of the maxilla and ectopterygoid, in the upper jaw. This functional modularity hypothesis is supported in the local superimposition but not common superimposition results, as the two functional modules are integrated together in the two most supported hypotheses of the common superimposition. The functional dissociation between the lateral and medial upper jaw was noted by Cundall (1983) because the maxilla plays a minimal direct role in ingestion. Yet, the morphological integration between the two suggested modules does not prevent the bones to be involved in different functions. Moreover, the maxilla articulates with the palatine and the ectopterygoid, which articulates with the pterygoid, and the presence of maxillary fangs dramatically restructures the morphology of the maxilla such that some ‘advanced’ (alethinophidian) snakes ingest prey using different mechanisms such as mandibular adduction because of the biomechanical limitations from modified cranial morphology due to the presence of maxillary fangs (Deufel and Cundall 2003). A modification in the morphology of the maxilla may then necessarily correspond with modifications in the other elements of the upper jaw because of these articulations, promoting integration within the upper jaw. This is especially true when considering positional information as in the common superimposition but is also captured in PLS1 of the maxilla-palatine integration, which shows that shape covariation between the two bones is dominated by their common joint surfaces (Fig. 4). So, while the upper jaw may behave as an evolutionary module, the maxilla and ectopterygoid’s incorporation into this module may be because of structural and physical associations with the palatopterygoid arch, rather than purely functional relationships. This is an important point because morphological integration caused by structural associations between bones is certainly present in the feeding system of snakes, however particularly structurally integrated, fused structures such as the avian cranium show highly complex patterns of modularity (seven modules, Felice & Goswami 2018) despite morphological evolution in one bone necessarily involving the evolution of neighboring bones because they must fit together. Yet, in the hyperkinetic snake feeding apparatus, the most supported hypotheses of modularity describe two to four modules out of eight completely unfused bones.

### Modularity and superimposition procedures

Comparing shape (co)variation of a common superimposition and local superimpositions enabled us to analyze separate mobile elements simultaneously and compare the strengths and weaknesses of each method for analyzing modularity. The presence of the maxilla in different modules depending on which superimposition method is used demonstrates the influence that incorporating positional information (GPA_all_) or isolating pure shape variation (local superimpositions) has when examining patterns of morphological modularity. The maxilla’s integration with the suspensorium when only considering pure shape variation (local superimpositions) is corroborated in PLS1 shape variation of the dentary, quadrate, maxilla, and ectopterygoid, which all showed consistent patterns of shape covariation with one another such that a robust dentary and maxilla, a quadrate with a wider proximal end, and a slender ectopterygoid lied at similar ends of PLS1, as did the combination of a slender dentary and maxilla, a slender and longer quadrate, and a wider anterior ectopterygoid (Fig. 4). This collection of bones that covary along consistent axes of shape covariation with each other may reflect a functional tradeoff between snakes adapted for elusive versus hard-bodied prey; a robust dentary and maxilla is better suited for prey capture than manipulation or ingestion, and a wider proximal quadrate provides additional area for attachment of the mandibular adductor muscle which may increase bite force (Fig. 1A). Comparing superimposition methods revealed competing factors contributing to the maxilla’s integration within the feeding system: local superimpositions revealed shared functional selective pressures integrating the maxilla with the suspensorium, and common superimposition revealed structural associations integrating the maxilla with the palatopterygoid arch. The fact that the common superimposition dataset showed a higher magnitude of overall morphological integration than the local superimposition dataset suggests that incorporating positional information contributed to the magnitude of integration (Supplementary Table 1). Allometry also had a greater effect on the common superimposition dataset, possibly due to backwards rotation of the quadrate dominating variation in landmark coordinates.

Although the different superimposition methods do not support the same first hypothesis of modularity, both methods strongly support Hypothesis 15 (Fig. 5, 6), describing four modules with relatively distinct functional roles. In this hypothesis of modularity, the palatopterygoid arch including the ectopterygoid form a module, driven by the highly conserved translational movement of the palatopterygoid arch during the pterygoid walk. The maxilla evolves as an individual module, possibly due to the competing structural versus functional influences explained above. The mandible is another module, highly integrated via selection for mechanical advantage. The quadrate and supratemporal, which articulate the feeding apparatus with the braincase, form the last module in this hypothesis, most likely coupled by a shared selective pressure for gape size. While the mandible also contributes to gape size, it is highly constrained by mechanical advantage as discussed above, and the backwards rotation of the quadrate dominates PC1 of the common superimposition (describing nearly half of the total shape variation, Fig. 1B), which is plausibly how, more specifically, a larger gape size is accomplished, thus making the quadrate and supratemporal its own evolutionary module. Further, the largest degree of evolutionary allometry was found in the quadrate; Palci and colleagues (2020) did not find significant evolutionary allometry in the quadrate when considering all Squamata, so the allometry found here likely relates to gape size and is adaptive.

Since different integration-inducing functional, developmental, and genetic processes may affect shape, positional, and size variation unevenly, our use of common and local superimpositions attempts to bracket the ‘true’ pattern of modularity. Moreover, eigenvalue dispersion and 2BPLS results, which are independent of rotational and positional variation, were largely consistent with CR results. H15 is possibly the pattern in which component parts of the feeding system evolve semi-independently, as both superimposition methods strongly support it, however there may exist an even less complex pattern of modularity considering that two-module hypotheses were most supported in both methods.

### Does the magnitude of morphological integration constrain morphological diversity?

Previous work has shown that the strength of integration can facilitate (Navalon et al. 2020; Fabre et al. 2020; Randau & Goswami 2017a), constrain (Goswami & Polly 2010b; Felice et al. 2018), or have no recoverable effect on morphological diversity (paedomorphic salamanders in Fabre et al. 2020; Bardua et al. 2019b; Watanabe et al. 2019; Bon et al. 2020). Here, we find that neither the strength of within-bone integration nor the average strength of each bone’s association with another has a significant effect on morphological diversity (Fig. 3B). As such, the considerably high degree of integration within the feeding system of aquatic-foraging snakes does not seem to affect morphological diversity over macroevolutionary timescales. This finding is interesting when considering the ecological and functional diversity of aquatic-foraging snakes, as it indicates that sufficient mechanical solutions to a broad range of feeding behaviors and diets are readily accessible within the highly integrated hyperkinetic feeding system of aquatic-foraging snakes.

### The hyperkinetic feeding system is highly integrated

Despite the kinesis in the feeding system of snakes, the individual bones are highly integrated with one another and organize into two to four evolutionary modules (Fig. 5, 6). In fact, the extreme kinesis is the reason why the feeding systems of snakes is so functionally optimized and thus so highly integrated (Cundall and Greene 2000; Moon et al. 2019). The developmental disintegration necessary to anatomically liberate fused structures implies that snake feeding bones oppose an expectation in morphological integration and modularity literature which suggests that functional systems may adaptively evolve in congruence with developmental systems or vice versa (Cheverud 1984, 1996; Wagner & Altenberg 1996; Klingenberg 2014). Of course, this does not consider alternative modes of developmental integration between separate bones, such as shared gene expressions, pleiotropic effects, or shared embryonic origin. Developmental systems can integrate spatially separated features as well, such as serially homologous limb bones (Hallgrimsson et al. 2002; Bell et al. 2011) or vertebrae (Randau & Goswami 2017a, b, 2018; Jones et al. 2018, 2020; Arlegi et al. 2020). Accordingly, it is possible that the evolutionary and functional modules of H15 (Fig. 5, 6) also match some pattern of developmental or genetic modularity. In any case, our developmental modularity hypotheses, which considered the embryonic origins of the articular and quadrate (splanchnocranium) and the rest of the feeding bones (dermatocranium), were not well supported (Supplementary Table 5). While additional patterns of developmental integration potentially contributing to morphological integration in the snake skull are less understood but surely exist, the high amount morphological integration in the kinetic feeding system of snakes is striking when considering the complex patterns of modularity recovered from the synostotic bones that make up akinetic morphological structures in mammals (Goswami 2006; Martín-Serra et al. 2018; Adams & Collyer 2019), archosaurs (Felice and Goswami 2018; Felice et al. 2019), and amphibians (Marshall et al. 2019; Bardua et al. 2019; Bon et al. 2020; Fabre et al. 2020). In the feeding system of aquatic-foraging snakes, we recognize strong functional and evolutionary integration generated because of its kinesis and developmental disintegration.

## Conclusion

In this paper, we quantify shape variation in the hyperkinetic feeding system of a phylogenetically broad sample of aquatic-foraging snakes and review patterns of morphological integration and modularity within this system. We find that the feeding system is highly integrated, with the most supported hypotheses of modularity involving only two modules despite there being eight separate bones unfused with one another. The most supported patterns of modularity describe an integrated palatopterygoid arch and mandible as separate modules, with the maxilla and quadrate either part of the palatopterygoid arch or mandible module depending on whether positional information was preserved in the superimposition method. Regardless, both superimposition methods show strong support for a four-module hypothesis with each separate module responsible for a specific functional role. This four-module hypothesis may be the best representation of how different regions of the feeding system independently evolve. Indeed, the major axes of phylogenetic-corrected shape variation of each of these modules have considerable functional consequences (Fig. 6), suggesting that modularity and integration is primarily influenced by performance-based selective pressures associated with feeding ecology. The utility of comparing common and local superimpositions of a mobile system to ‘bracket’ the most biologically accurate pattern of modularity proved fruitful and may be considered in future studies when taking appropriate precautions. Further, the relatively high degree of integration in this hyperkinetic system is fascinating when considering the developmental disintegration necessary to spatially disintegrate each component part, and the complex patterns of modularity found in fused structures such as rodent mandibles (Adams & Collyer 2019). Despite this exceptionally strong integration within the feeding system, morphological diversity is not apparently constrained, indicating that adequate mechanical and functional solutions to a wide variety of dietary and ecological niches are readily available within constraint due to integration in the feeding system. Further research addressing morphological integration in the skull of snakes with different dietary challenges (e.g. arboreal species, snail-eaters, egg-eaters) should be conducted to reveal the relative importance of competing functional, developmental, and genetic factors influencing morphological integration and their micro- and macroevolutionary consequences.

## Acknowledgments

We thank the herpetological collections staff of the American Museum of Natural History: David Kizirian, David Dickey, Margaret Arnold and especially Lauren Vonnahme, but also Alan Resetar (Field Museum of Natural History), Erica Ely and Lauren Scheinberg (California Academy of Sciences) for their help and patience in carefully choosing specimens that fit our study and quickly processing specimen loans. Another special thanks to Morgan Hill Chase and Andrew Smith, from the Microscopy and Imaging and Facility who did all the CT scanning involved in this study. The authors greatly appreciate insightful comments by Spencer Hellert that improved earlier versions of this manuscript. We thank the National Science Foundation REU program and the Fyssen Foundation for partly funding this research.

## Author contributions

DR contributed to the data acquisition, analysis, interpretation of the results, writing and editing. DP and CR contributed to the interpretation and discussion of the results, as well as the review and edits of the manuscript. MS conceived the study, generated the scans, helped in the analyses, the interpretation of results and reviewed the manuscript.

## Supplementary Tables & Figures

**Supplementary Table 1.**
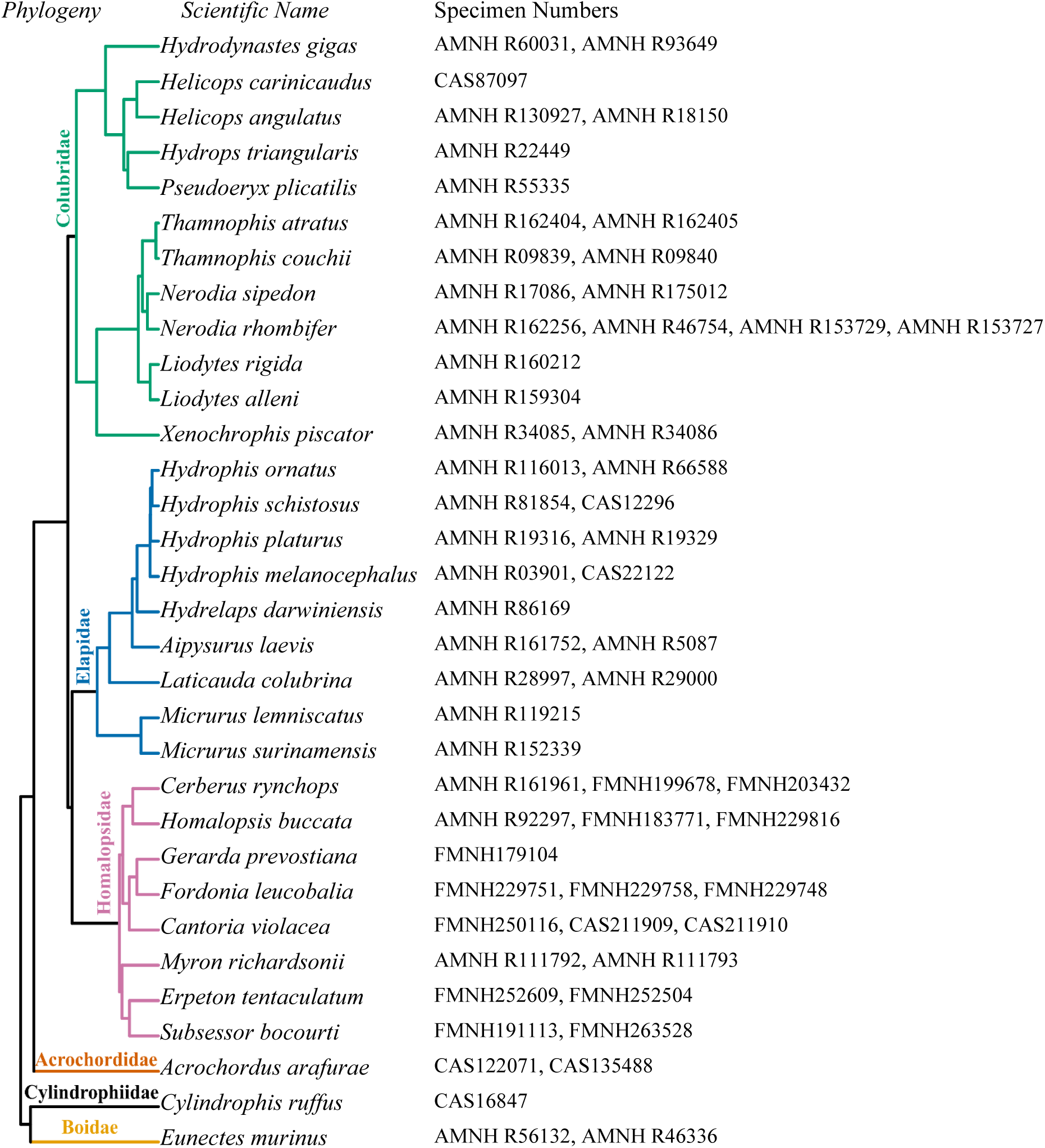
Species and specimens used in this study. Museum codes: AMNH – American Museum of Natural History, FMNH – Field Museum of Natural History, CAS – California Academy of Sciences. Topology of the phylogeny used in this paper, from Pyron & Burbrink (2014).

**Supplementary Table 2.** Explanations for each alternative hypothesis of modularity.

| # | Hypothesis | Explanation |
| --- | --- | --- |
| 1 | [dentary, compound] + [else] | Cranial elements and mandibular elements as separate evolutionary modules. Dorsal-ventral modularity. |
| 2 | [dentary, compound, quadrate, supratemporal] + [pterygoid, palatine, ectopterygoid, maxilla] | Suspensorium module consisting of the mandible, quadrate, and supratemporal and an upper jaw module. The mandible, quadrate, and supratemporal cumulatively contribute to gape size, which dictates the maximum size of prey (Cundall & Greene 2000). The elements of the suspensorium must act in concert to successfully strike at prey (Cundall & Greene 2000; Moon et al. 2019). |
| 3 | [maxilla, ectopterygoid] + [else] | The maxilla is not functionally integrated as strongly as the other bones (Cundall 1983), however the presence of fangs restructures its morphology and function (Kardong 1979; Vonk et al. 2008), and it may be integrated with the ectopterygoid because they articulate. |
| 4 | [dentary, compound, quadrate, maxilla] + [supratemporal, pterygoid, palatine, ectopterygoid] | Lateral vs. medial modules |
| 5 | [dentary, anterior compound] + [else] | Same as hypothesis #1 but with developmental modularity in compound. The posterior compound and quadrate are splanchnocranial. |
| 6 | [pterygoid, palatine] + [else] | Two-module hypothesis with the palatopterygoid arch as an individual module. The pterygoid and palatine articulate to make the palatopterygoid arch, which is necessary for the successfully functioning of the 'pterygoid walk' (Bolt & Ewer 1964). |
| 7 | [quadrate, supratemporal] + [else] | The quadrate's size and relative orientation varies greatly among species, and it is the only bone involved in prey detection as it connects to the stapes. The supratemporal is articulated with the quadrate and its relative position also varies greatly across the dataset. |
| 8 | [posterior compound, quadrate, supratemporal] + [else] | Developmental modularity; the quadrate and articular of the compound (posterior compound) are splanchnocranial, the other bones are dermatocranial. |
| 9 | [dentary, compound] + [quadrate, supratemporal] + [pterygoid, palatine, ectopterygoid, maxilla] | Same as hypothesis #1 and #2, but with the quadrate and supratemporal as an individual module. The relative orientation of the quadrate varies greatly among species, which has biomechanical implications (Scanferla 2016). |
| 10 | [dentary, compound, quadrate, supratemporal] + [maxilla] + [pterygoid, palatine, ectopterygoid] | Same as hypothesis #2, but with the mandible (suspensorium) as an individual module. |
| 11 | [dentary, compound, quadrate, supratemporal] + [maxilla, ectopterygoid] + [pterygoid, palatine] | Same as hypothesis #11, but with the ectopterygoid integrated with the maxilla, because they articulate. |
| 12 | [dentary, maxilla] + [compound, quadrate, supratemporal] + [pterygoid, palatine, ectopterygoid] | The dentary and maxilla are tooth bearing bones which are in direct physical contact with prey during prey capture (Cundall 1983; Cundall & Greene 2000; Moon et al. 2019). The compound and quadrate are connected via the mandibular adductor, which originates on the anterior quadrate and inserts on the posterolateral compound (Johnston 2014). The palatopterygoid arch articulates with the ectopterygoid. |
| 13 | [dentary, compound, maxilla, ectopterygoid] + [quadrate, supratemporal] + [pterygoid, palatine] | Same as hypothesis #11, but with the compound and ectopterygoid in the same module as the dentary and maxilla because they articulate with the dentary and maxilla, respectively. |
| 14 | [dentary, anterior compound] + [posterior compound, quadrate, supratemporal] + [pterygoid, palatine, ectopterygoid, maxilla] | Same as hypothesis #9 but incorporating developmental modularity in the compound. |
| 15 | [dentary, compound] + [quadrate, supratemporal] + [pterygoid, palatine, ectopterygoid] + [maxilla] | Same as hypothesis #9, but with the maxilla as an individual module. The maxilla has little direct role in any functional module other than prey capture in many species (Cundall 1983). |
| 16 | [dentary, compound] + [quadrate, supratemporal] + [pterygoid, palatine] + [maxilla, ectopterygoid] | Same as hypothesis #15, but with the ectopterygoid in the maxilla module. |
| 17 | [dentary, anterior compound] + [posterior compound, quadrate, supratemporal] + [pterygoid, palatine] + [ectopterygoid, maxilla] | Same as hypothesis #16, but modularity within compound. |
| 18 | Complete modularity except [pterygoid, palatine] + [dentary, compound] | The mandible and palatopterygoid arch are hypothesized to be two of the most integrated pairs of bones. This hypothesis considers modularity except for these two structures. |
| 19 | Complete modularity except [pterygoid, palatine] | Same as hypothesis #5, but with complete modularity outside of the palatopterygoid arch. |
| 20 | Complete modularity except [dentary, compound] | The mandibular elements are expected to be integrated because of their articulation and functional relationships; are mandibular, but not cranial elements integrated? |
| 21 | Complete modularity | The separate bones are unfused with one another, potentially allowing one bone to evolve without the necessary evolution of another. Does cranial kinesis promote modularity? |

**Supplementary Table 3.** The influence of allometry, phylogeny, and integration on snake skull bones associated with feeding. Results of allometry (PGLS), phylogenetic signal (all significant; 0.001 < p < 0.004), and eigenvalue dispersion analyses. Significant p-values of allometry analyses are in bold. Higher eigenvalue dispersion value corresponds to higher degrees of overall integration within each structure. That is, more integrated structures will have more correlated shape variation consolidated in the first few principal components (eigenvalues).

| Structure | Allometry |  | Phylogenetic Signal |  | Integration |
| --- | --- | --- | --- | --- | --- |
|  | R<sup>2</sup> | P | K | Effect Size | Eigenvalue dispersion |
| Dentary | 0.06 | 0.15 | 0.45 | 2.99 | 0.61 |
| Compound | 0.08 | 0.051 | 0.58 | 3.91 | 0.65 |
| Quadrate | **0.26** | **0.001** | 0.51 | 3.05 | 0.55 |
| Supratemporal | **0.08** | **0.02** | 0.47 | 3.76 | 0.45 |
| Pterygoid | 0.07 | 0.09 | 0.51 | 3.04 | 0.47 |
| Ectopterygoid | 0.03 | 0.39 | 0.38 | 3.13 | 0.65 |
| Palatine | **0.08** | **0.02** | 0.59 | 5.97 | 0.48 |
| Maxilla | **0.07** | **0.03** | 0.57 | 4.63 | 0.47 |
| Common Superimposition | **0.13** | **0.001** | 0.48 | 4.20 | 0.45 |
| Local Superimpositions | **0.07** | **0.02** | 0.52 | 4.90 | 0.35 |

**Supplementary Table 4.** The r-PLS values of PLS1 (above diagonal) and p-values (below diagonal) of each 2BPLS analysis. Significant values are in bold.

|  | Dentary | Compound | Quadrate | Supratemporal | Pterygoid | Ectopterygoid | Palatine | Maxilla |
| --- | --- | --- | --- | --- | --- | --- | --- | --- |
| Dentary |  | **0.634** | **0.619** | **0.665** | 0.476 | **0.822** | 0.651 | **0.711** |
| Compound | **0.009** |  | 0.589 | 0.465 | 0.407 | **0.573** | 0.561 | 0.578 |
| Quadrate | **0.043** | 0.053 |  | **0.674** | **0.647** | **0.686** | 0.651 | **0.65** |
| Supratemporal | **0.029** | 0.49 | **0.013** |  | 0.560 | **0.73** | **0.692** | **0.686** |
| Pterygoid | 0.158 | 0.312 | **0.022** | 0.111 |  | 0.453 | **0.74** | 0.606 |
| Ectopterygoid | **0.001** | **0.043** | **0.028** | **0.012** | 0.197 |  | 0.611 | **0.806** |
| Palatine | 0.051 | 0.123 | 0.051 | **0.02** | **0.003** | 0.105 |  | **0.747** |
| Maxilla | **0.009** | 0.115 | **0.045** | **0.041** | 0.087 | **0.001** | **0.003** |  |

**Supplementary Table 5.** The z-PLS values of PLS1 (above diagonal) and p-values (below diagonal) of each 2BPLS analysis. Significant values are in bold.

|  | Dentary | Compound | Quadrate | Supratemporal | Pterygoid | Ectopterygoid | Palatine | Maxilla |
| --- | --- | --- | --- | --- | --- | --- | --- | --- |
| Dentary |  | **2.905** | **1.910** | **1.980** | 1.035 | **4.058** | 1.818 | **2.435** |
| Compound | **0.009** |  | 1.827 | 0 | 0.389 | **2.073** | 1.192 | 1.262 |
| Quadrate | **0.043** | 0.053 |  | **2.362** | **2.342** | **2.348** | 1.852 | **1.874** |
| Supratemporal | **0.029** | 0.49 | **0.013** |  | 1.255 | **2.512** | **2.138** | **1.766** |
| Pterygoid | 0.158 | 0.312 | **0.022** | 0.111 |  | 0.8233 | **3.040** | 1.452 |
| Ectopterygoid | **0.001** | **0.043** | **0.028** | **0.012** | 0.197 |  | 1.4378 | **3.272** |
| Palatine | 0.051 | 0.123 | 0.051 | **0.02** | **0.003** | 0.105 |  | **2.944** |
| Maxilla | **0.009** | 0.115 | **0.045** | **0.041** | 0.087 | **0.001** | **0.003** |  |

**Supplementary Table 6.** Alternative hypotheses of modularity and their support, measured as phylogenetic-corrected Z_CR_, for both common and local superimposition procedures. The lower the Z_CR_ and CR values, the stronger the modular signal. ‘Else’ refers to all the other bones not yet mentioned as part of their own module. Rows are ordered by Z_CR_ of common superimposition and are colored according to strength of modular signal.

| # | Hypothesis | Modules | Common Superimposition |  | Local Superimpositions |  |
| --- | --- | --- | --- | --- | --- | --- |
| | | | CR | $Z_{CR}$ | CR | $Z_{CR}$ |
| 1 | [dentary, compound] + [else] | 2 | 0.63 | -31.18 | 0.65 | -27.02 |
| 2 | [dentary, compound, quadrate, supratemporal] + [pterygoid, palatine, ectopterygoid, maxilla] | 2 | 0.76 | -31.03 | 0.77 | -26.08 |
| 15 | [dentary, compound] + [quadrate, supratemporal] + [pterygoid, palatine, ectopterygoid] + [maxilla] | 4 | 0.72 | -30.93 | 0.62 | -28.13 |
| 9 | [dentary, compound] + [quadrate, supratemporal] + [pterygoid, palatine, ectopterygoid, maxilla] | 3 | 0.76 | -30.86 | 0.74 | -25.52 |
| 16 | [dentary, compound] + [quadrate, supratemporal] + [pterygoid, palatine] + [maxilla, ectopterygoid] | 4 | 0.76 | -30.75 | 0.76 | -24.79 |
| 10 | [dentary, compound, quadrate, supratemporal] + [maxilla] + [pterygoid, palatine, ectopterygoid] | 3 | 0.74 | -30.68 | 0.58 | -28.12 |
| 17 | [dentary, anterior compound] + [posterior compound, quadrate, supratemporal] + [pterygoid, palatine] + [ectopterygoid, maxilla] | 4 | 0.77 | -30.61 | 0.76 | -24.50 |
| 4 | [dentary, compound, quadrate, maxilla] + [supratemporal, pterygoid, palatine, ectopterygoid] | 2 | 0.82 | -30.54 | 0.68 | -28.14 |
| 12 | [dentary, maxilla] + [compound, quadrate, supratemporal] + [pterygoid, palatine, ectopterygoid] | 3 | 0.81 | -30.53 | 0.73 | -26.69 |
| 14 | [dentary, anterior compound] + [posterior compound, quadrate, supratemporal] + [pterygoid, palatine, ectopterygoid, maxilla] | 3 | 0.78 | -30.47 | 0.77 | -24.24 |
| 19 | Complete modularity except [pterygoid, palatine] | 7 | 0.71 | -30.37 | 0.65 | -24.35 |
| 21 | Complete modularity | 8 | 0.73 | -30.21 | 0.67 | -24.72 |
| 11 | [dentary, compound, quadrate, supratemporal] + [maxilla, ectopterygoid] + [pterygoid, palatine] | 3 | 0.81 | -30.15 | 0.81 | -21.25 |
| 5 | [dentary, anterior compound] + [else] | 2 | 0.74 | -30.11 | 0.74 | -22.45 |
| 18 | Complete modularity except [pterygoid, palatine] + [dentary, compound] | 6 | 0.75 | -29.78 | 0.68 | -23.66 |
| 20 | Complete modularity except [dentary, compound] | 7 | 0.77 | -29.42 | 0.72 | -22.47 |
| 3 | [maxilla, ectopterygoid] + [else] | 2 | 0.85 | -29.13 | 0.85 | -18.27 |
| 8 | [posterior compound, quadrate, supratemporal] + [else] | 2 | 0.90 | -28.12 | 0.98 | -3.57 |
| 6 | [pterygoid, palatine] + [else] | 2 | 0.90 | -25.91 | 0.83 | -17.53 |
| 13 | [dentary, compound, maxilla, ectopterygoid] + [quadrate, supratemporal] + [pterygoid, palatine] | 3 | 0.92 | -25.26 | 0.86 | -18.04 |
| 7 | [quadrate, supratemporal] + [else] | 2 | 0.98 | -12.00 | 0.98 | -3.36 |

**Supplemental Figure 1.**
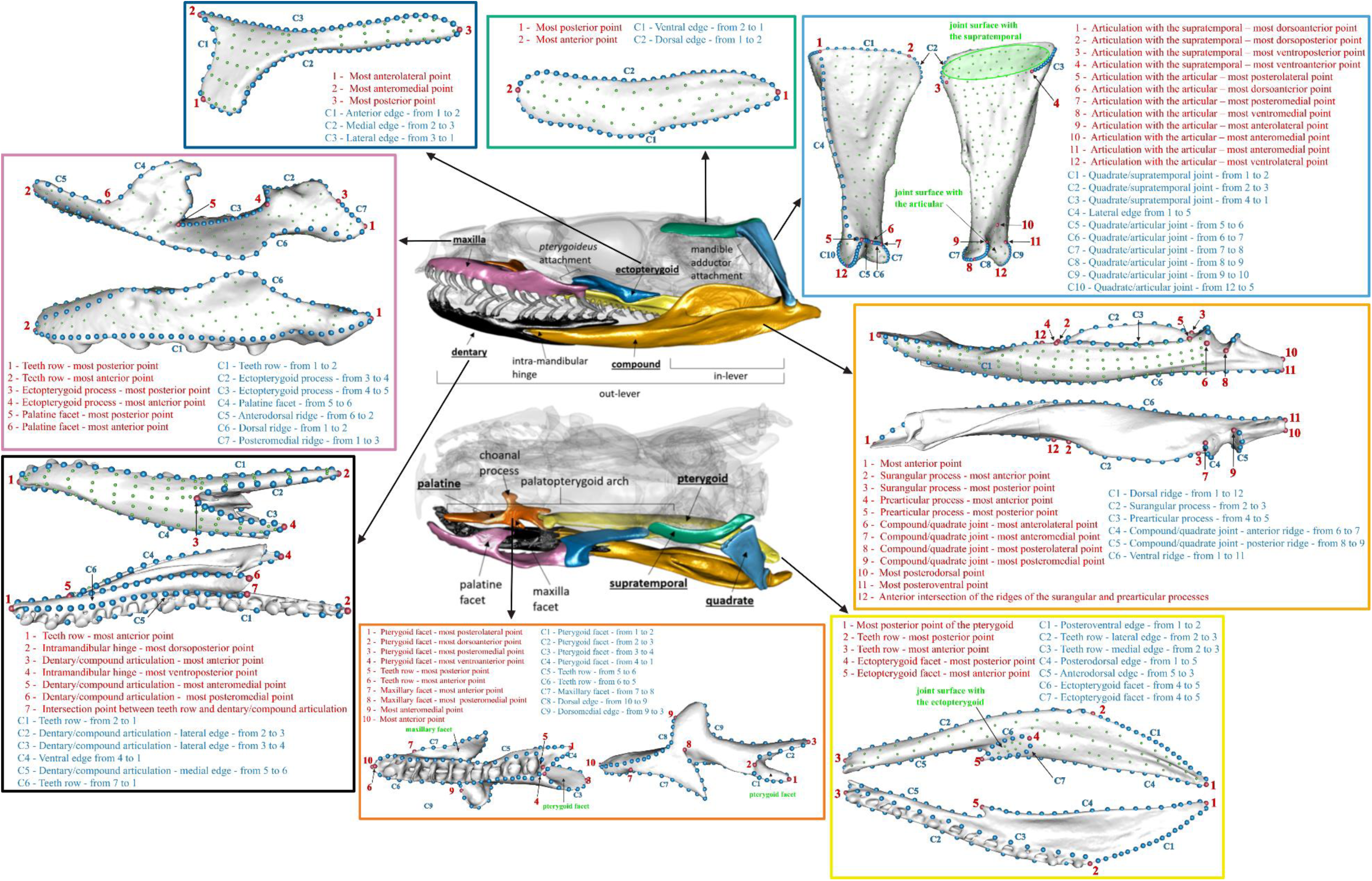
The bones analyzed in this paper (middle two skulls) and their respective landmark configurations. Red indicates anatomical landmarks, blue curve semi-landmarks and green surface semi-landmarks, along with a list of description of each landmark (red) and curves (blue) and some important anatomical features (green).

**Supplemental Figure 2.**
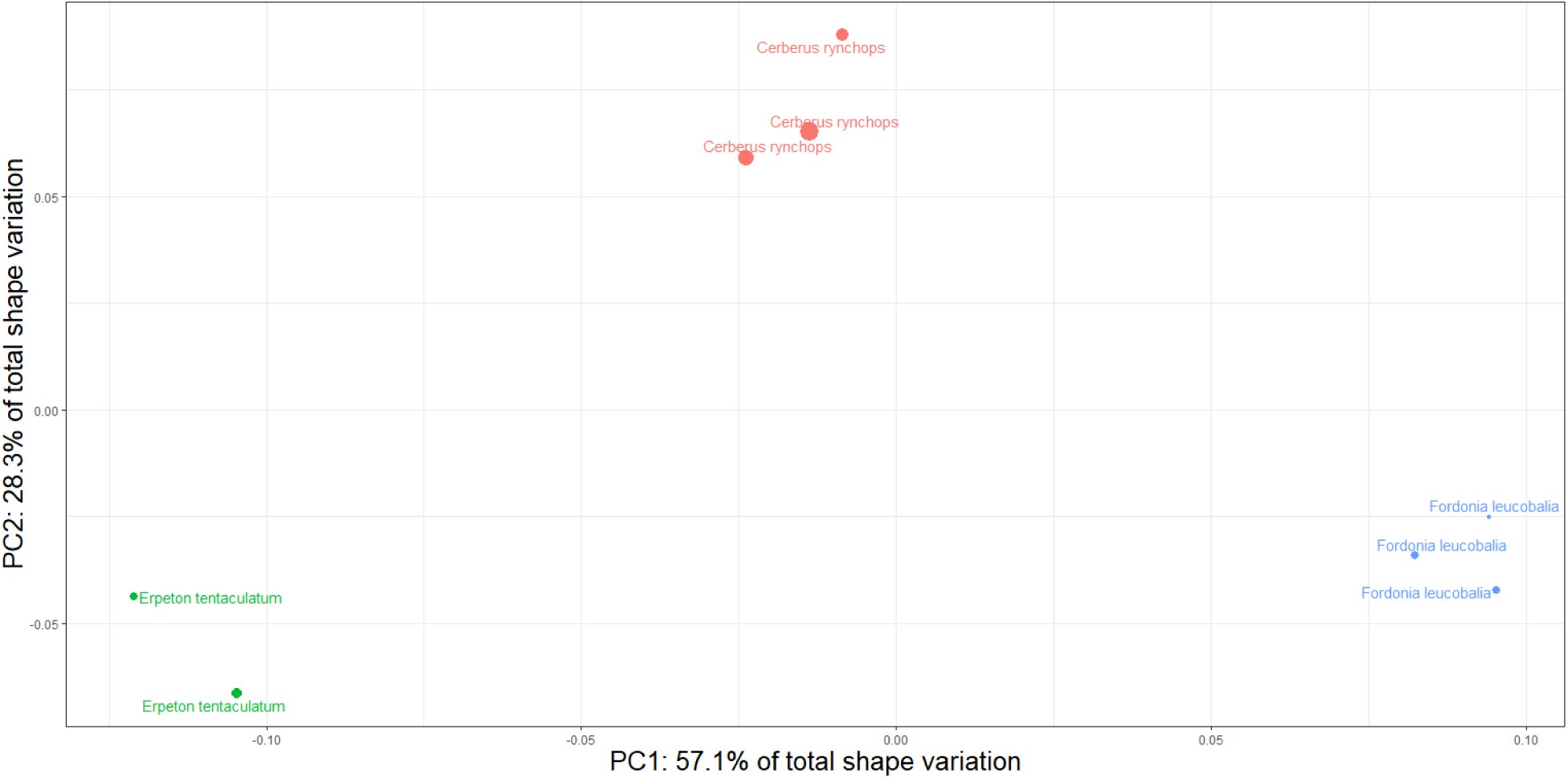
Principal component morphospace from a common superimposition (GPA_All_) of individual specimens of three homalopsid snakes: *Erpeton tentaculatum* (green), *Fordonia leucobalia* (blue), and *Cerberus rynchops* (orange). Note that individual specimens of each species group together much more closely than different species.

**Supplementary Figure 3.**
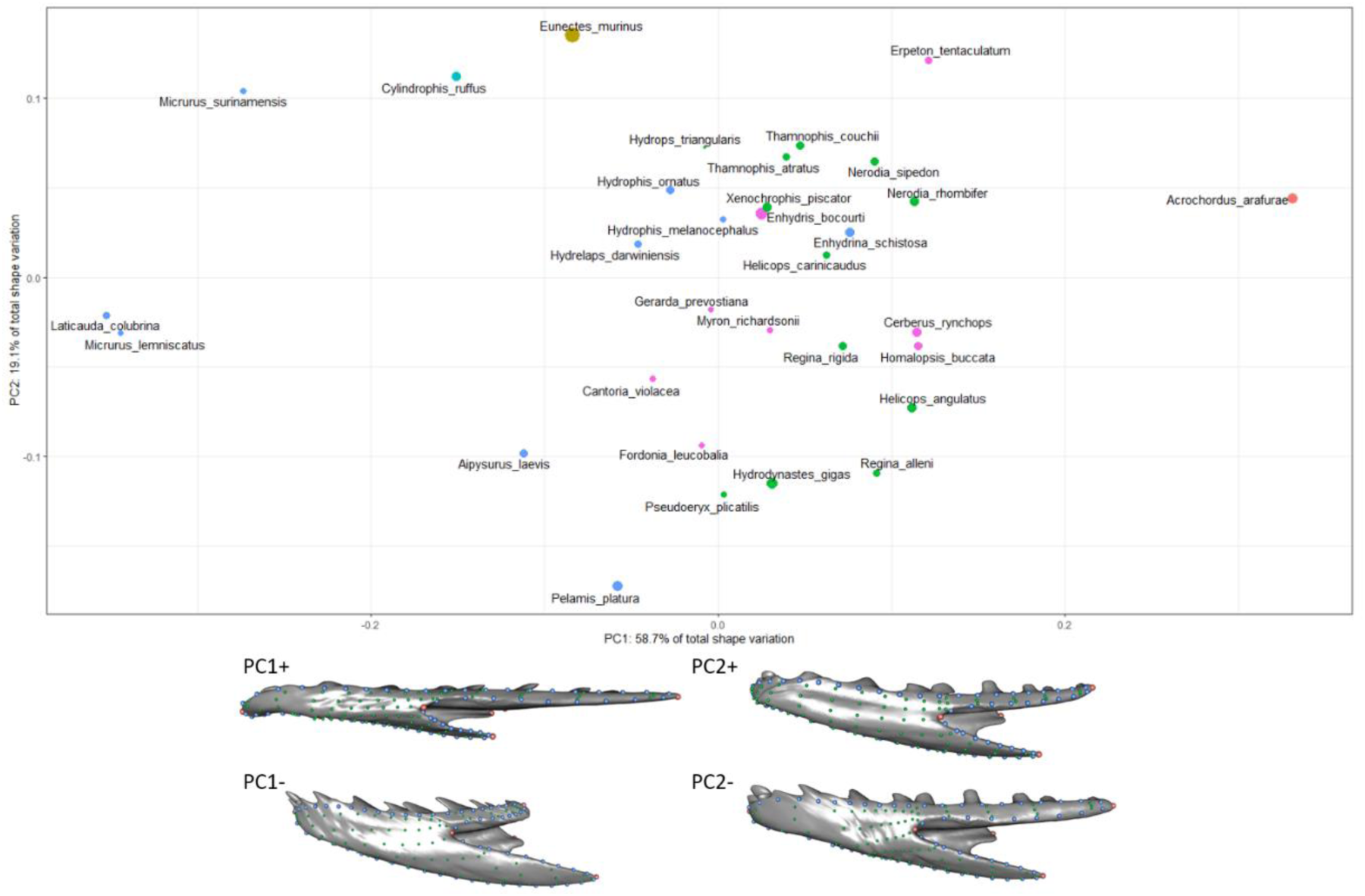
Morphospace of the first two principal components of the dentary (above), and axes of shape variation along the first two principal components in lateral view (below). Size of each point (species) corresponds to centroid size of the dentary and color corresponds to taxonomic family as in Figure 1. This scheme is consistent for Supplementary figures 3-10.

**Supplementary Figure 4.**
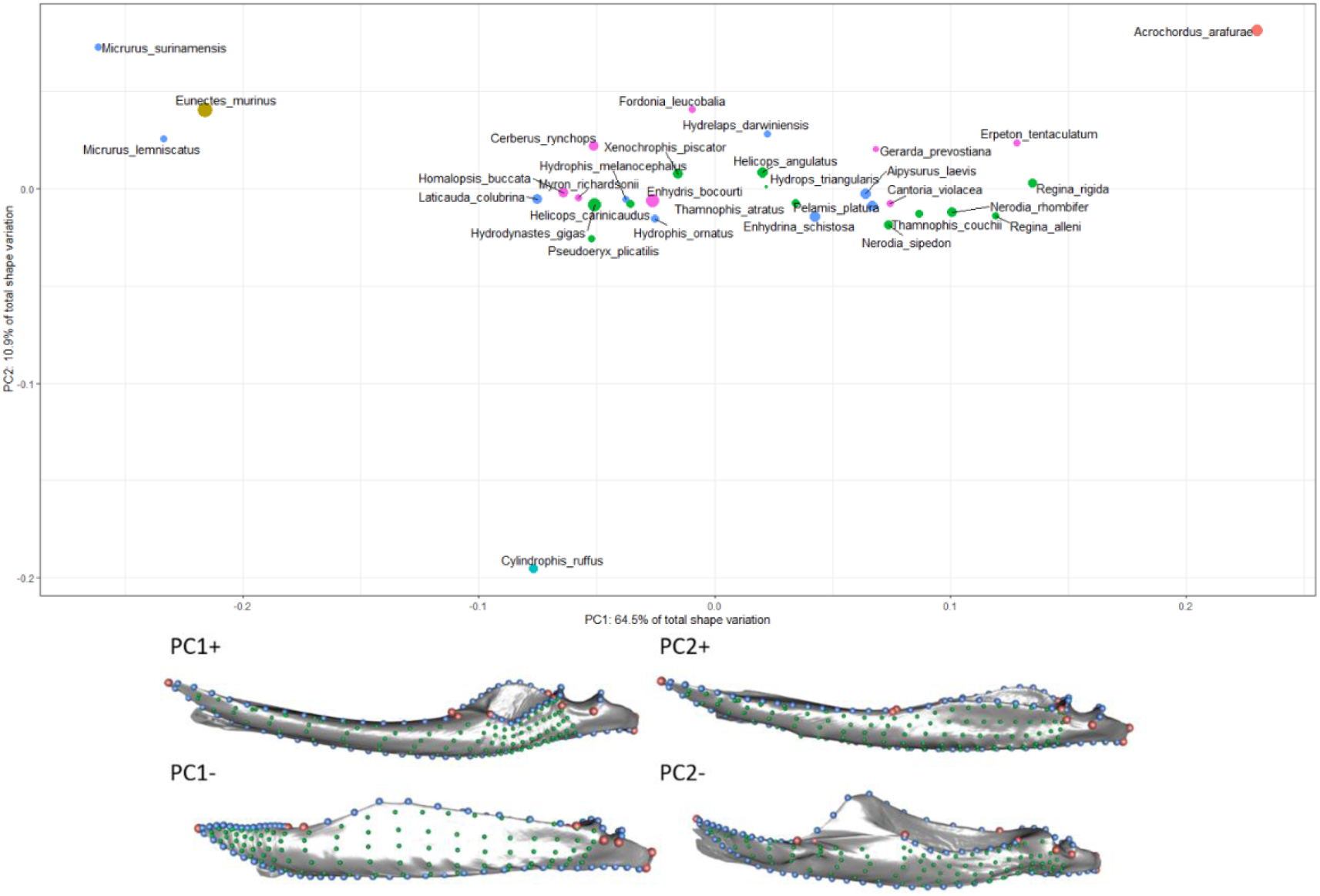
Morphospace of the first two principal components of the compound (above), and axes of shape variation along the first two principal components in lateral view (below).

**Supplementary Figure 5.**
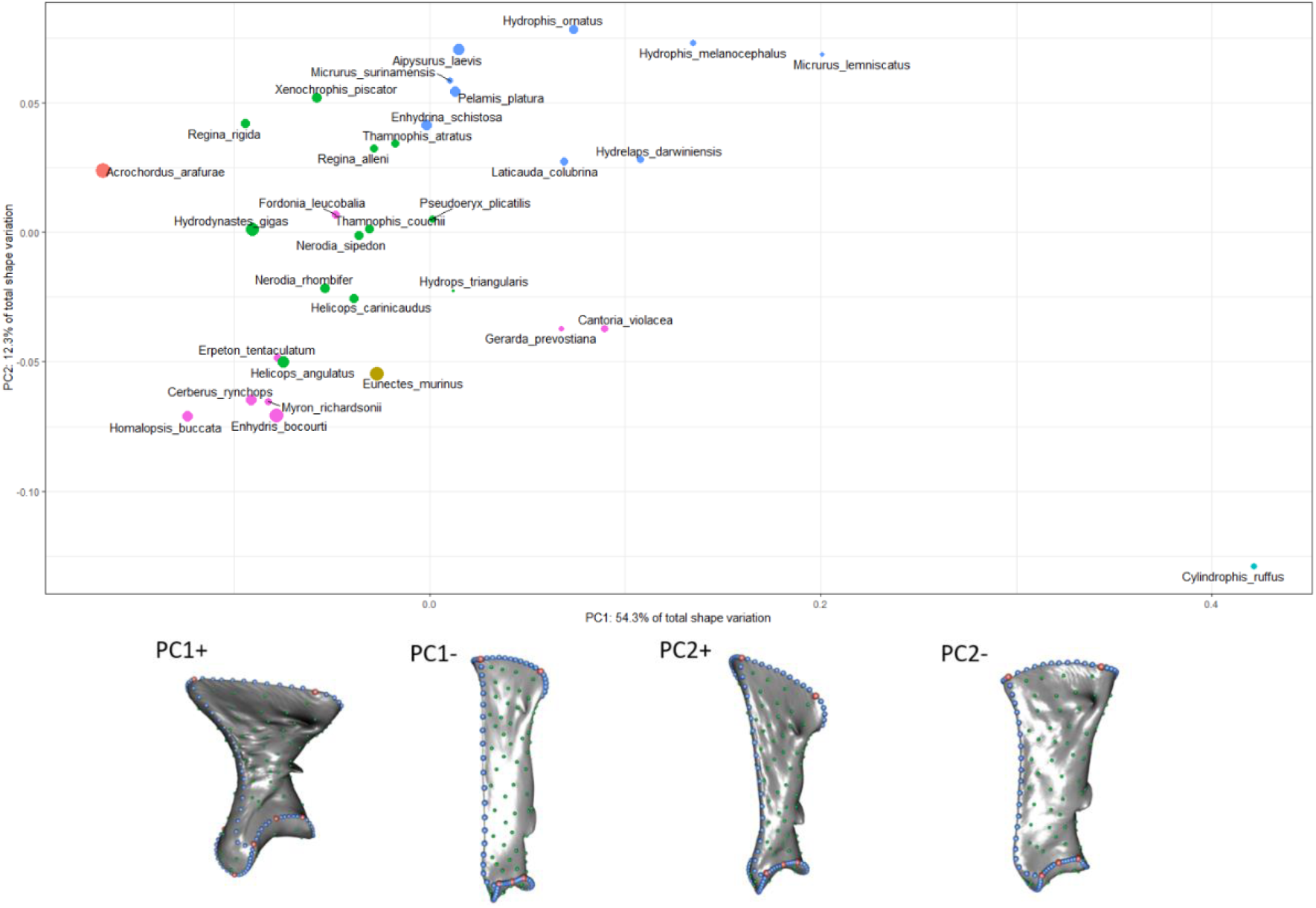
Morphospace of the first two principal components of the quadrate (above), and axes of shape variation along the first two principal components in lateral view (below).

**Supplementary Figure 6.**
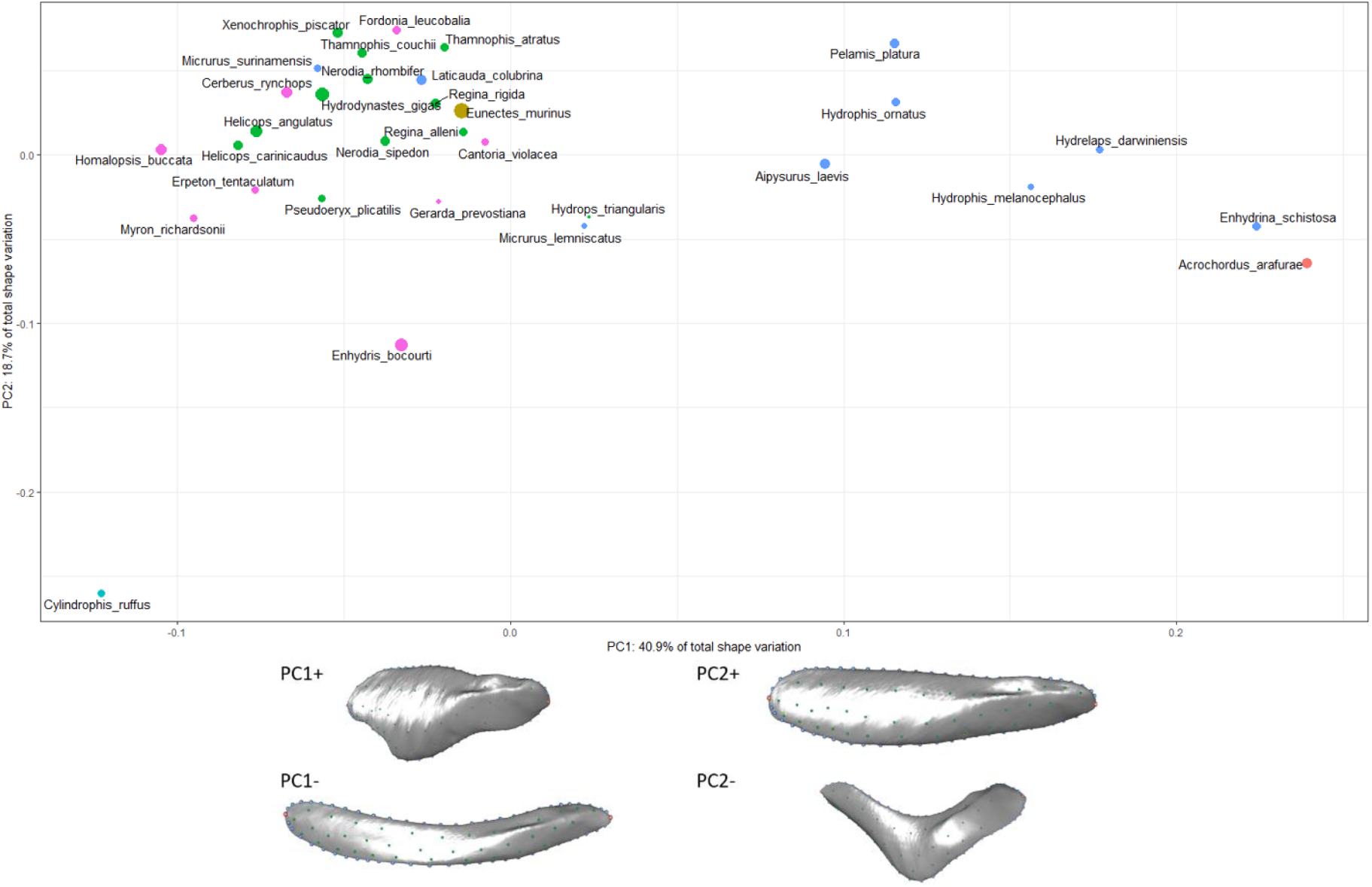
Morphospace of the first two principal components of the supratemporal (above), and axes of shape variation along the first two principal components in lateral view (below).

**Supplementary Figure 7.**
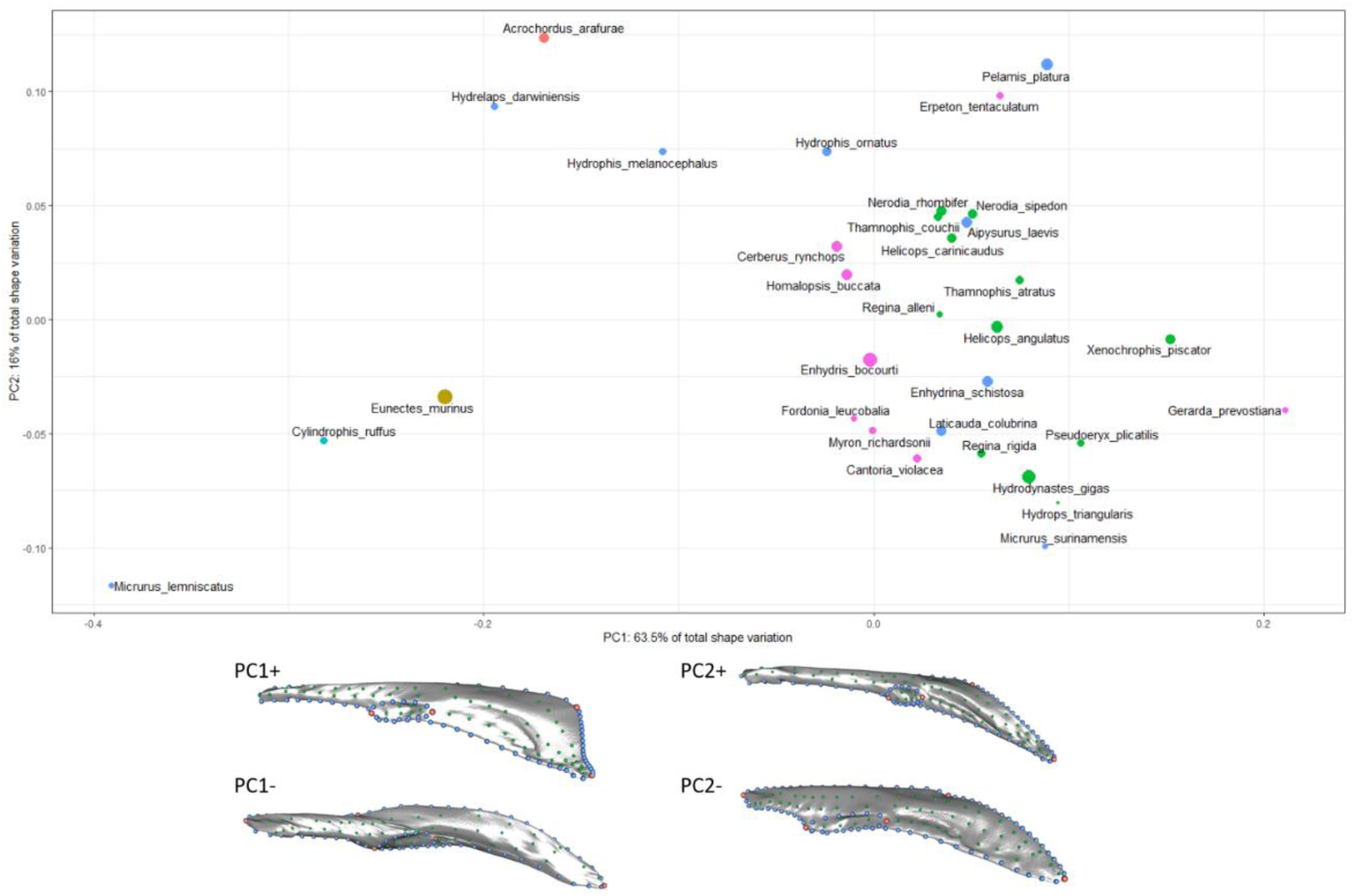
Morphospace of the first two principal components of the pterygoid (above), and axes of shape variation along the first two principal components in dorsal view (below).

**Supplementary Figure 8.**
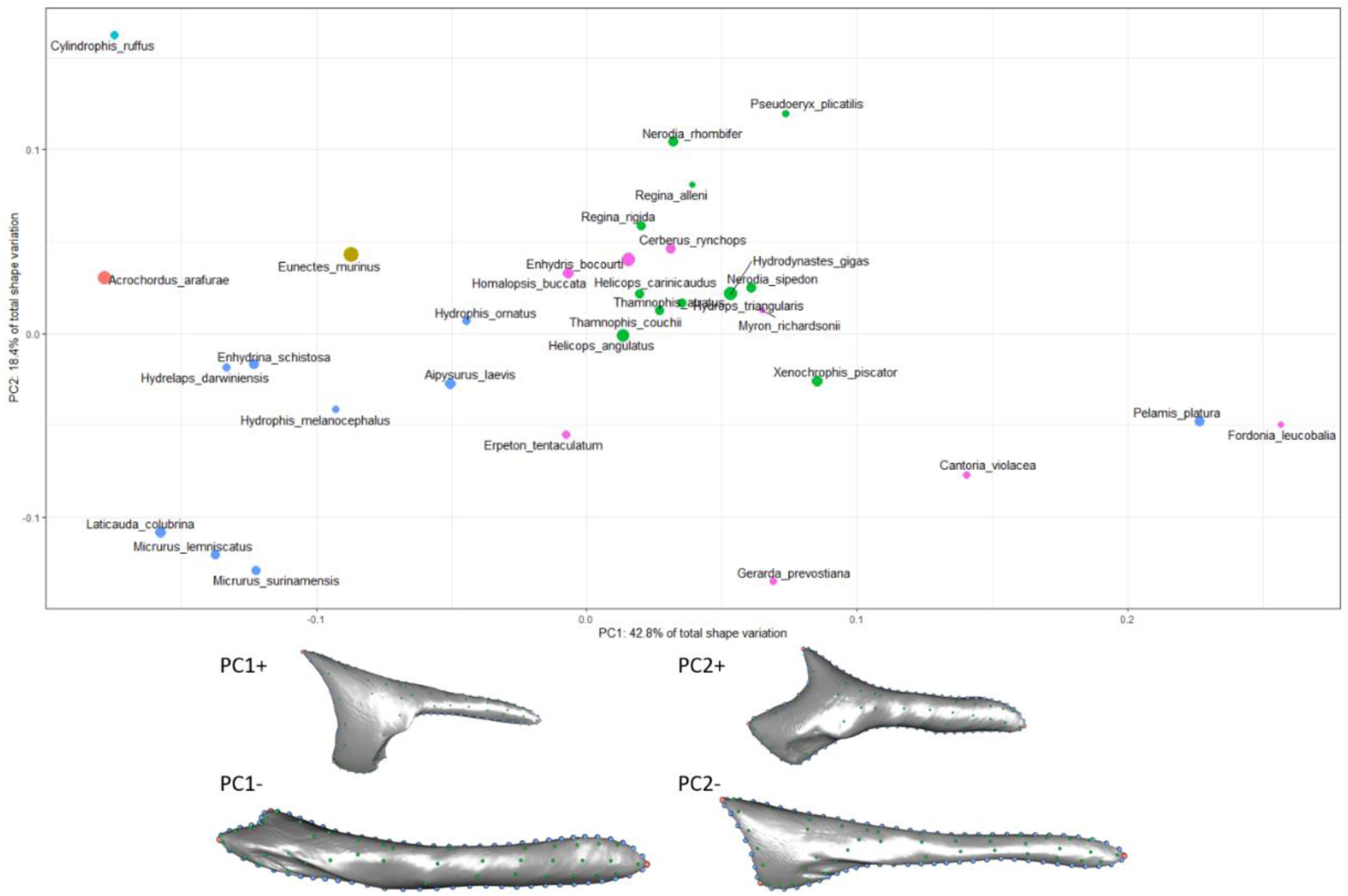
Morphospace of the first two principal components of the ectopterygoid (above), and axes of shape variation along the first two principal components in dorsal view (below).

**Supplementary Figure 9.**
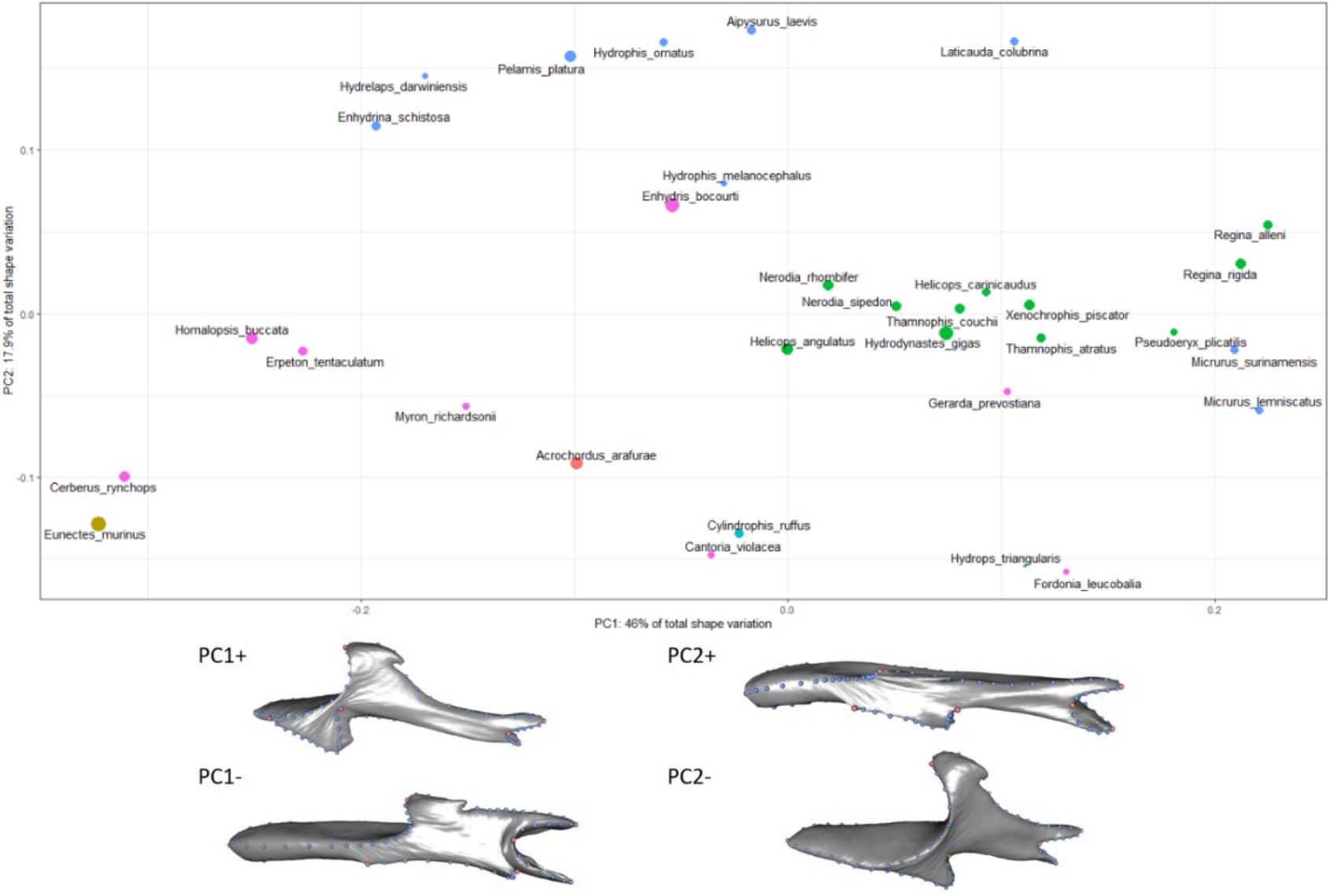
Morphospace of the first two principal components of the palatine (above), and axes of shape variation along the first two principal components in dorsal view (below).

**Supplementary Figure 10.**
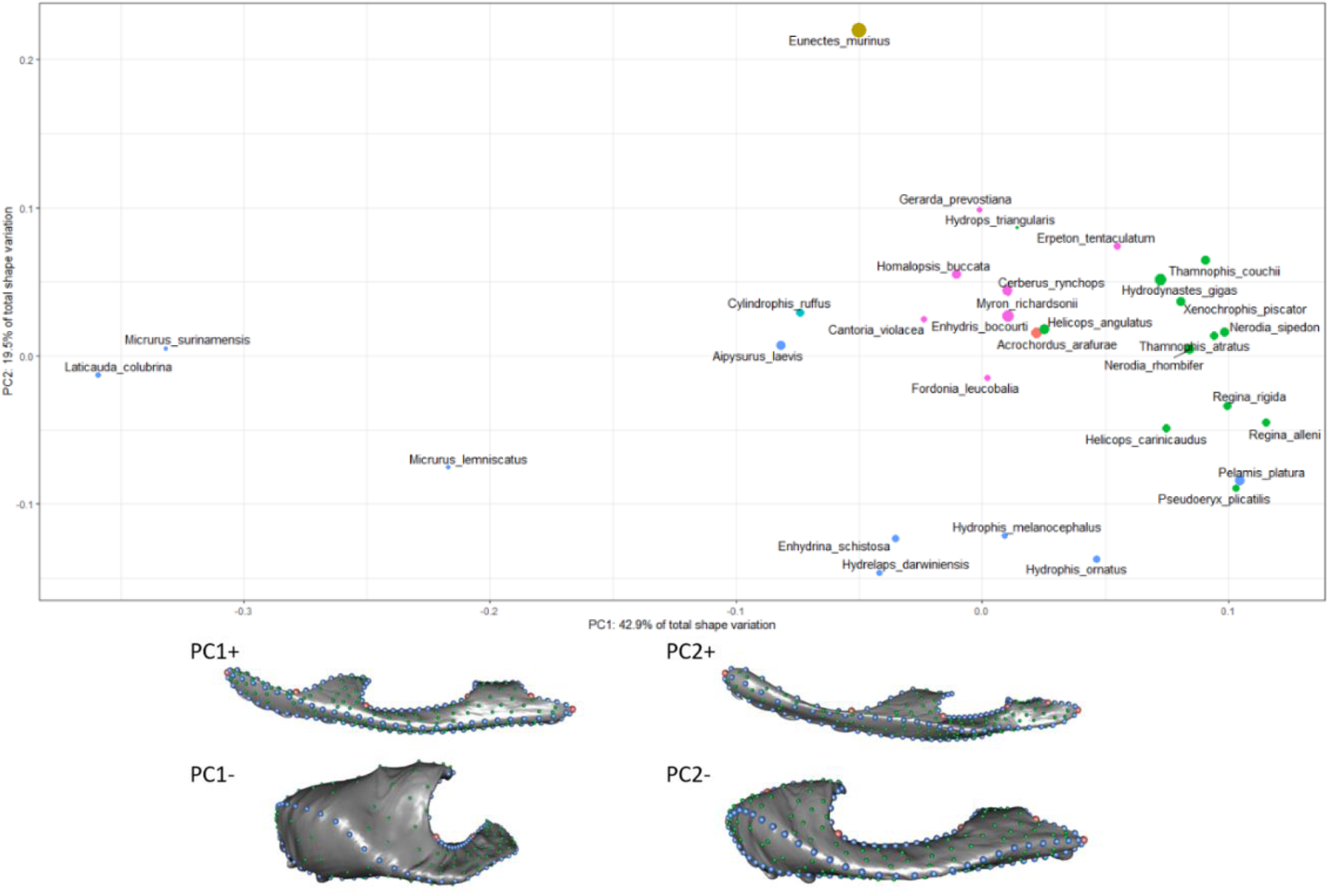
Morphospace of the first two principal components of the maxilla (above), and axes of shape variation along the first two principal components in dorsal view (below).

**Supplementary Figure 11.**
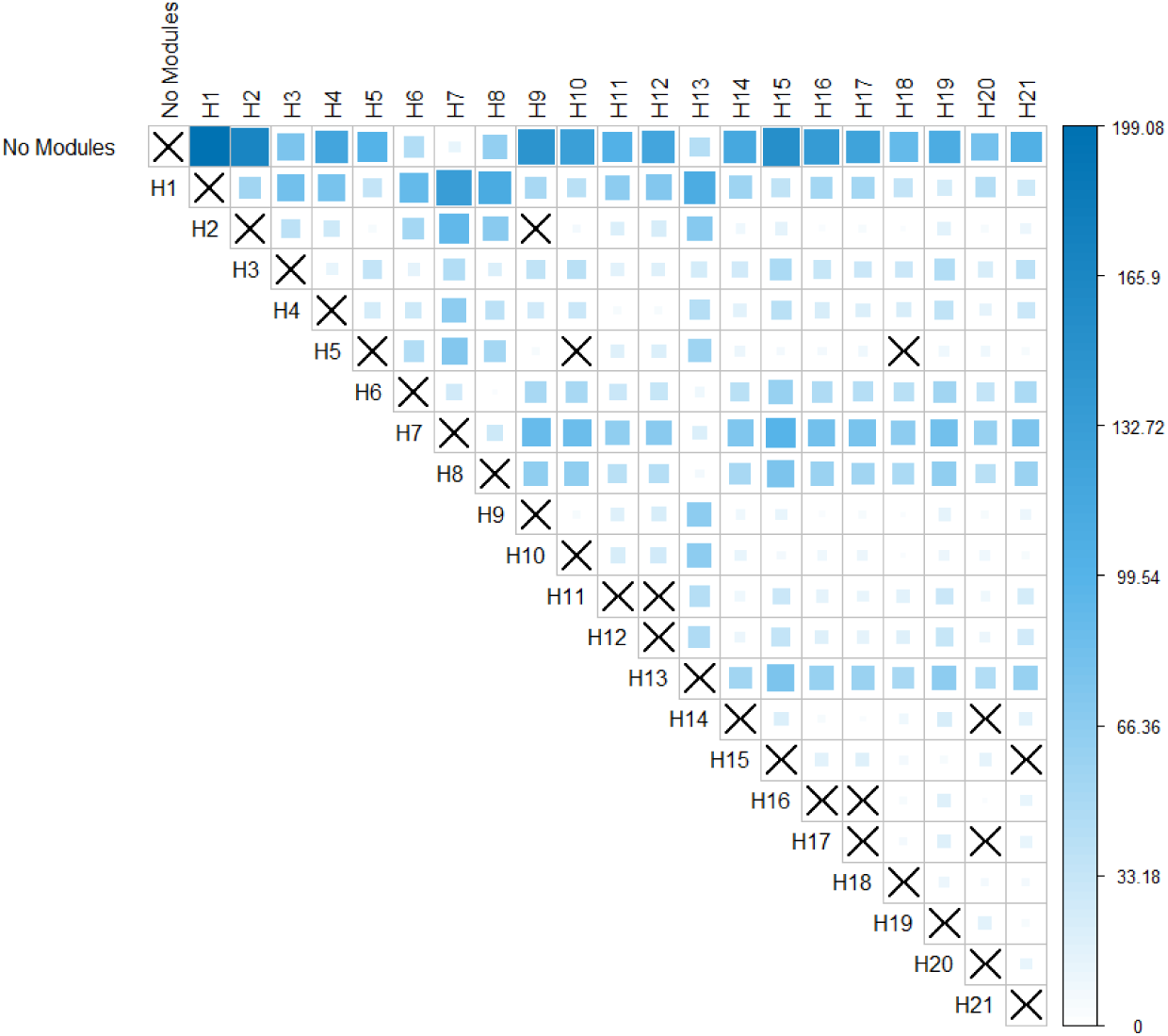
Pairwise effect sizes of alternative hypotheses of modularity of the common superimposition. Darker blue cells correspond to higher effect sizes and larger differences between strength of modular signal between alternative hypotheses. Insignificant (i.e., not significantly different support) values are marked by ‘X’s. ‘No Modules’ refers to the null hypothesis of complete integration.

**Supplementary Figure 12.**
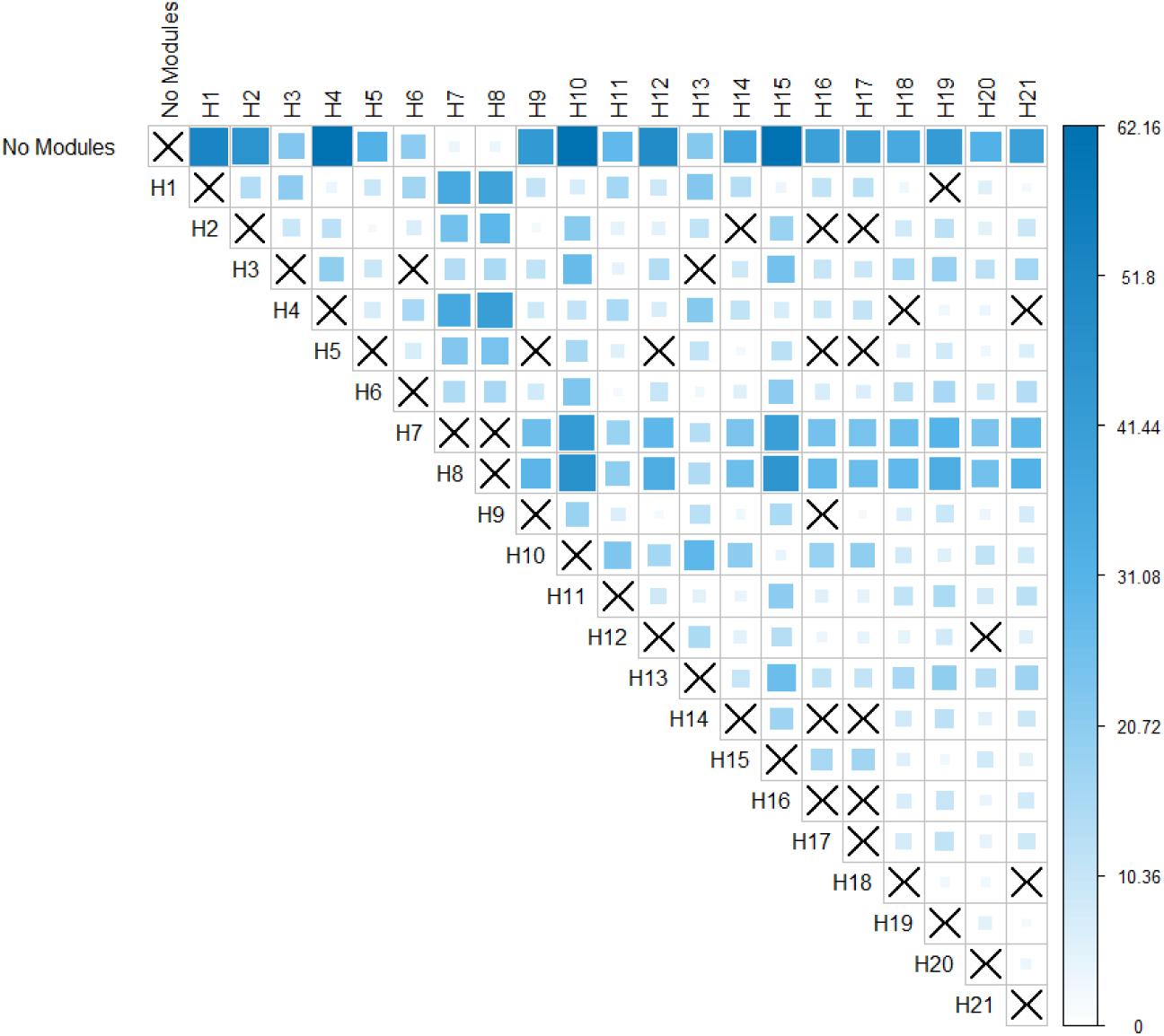
Pairwise effect sizes of alternative hypotheses of modularity of the local superimposition dataset. Darker blue cells correspond to higher effect sizes and larger differences between strength of modular signal between alternative hypotheses. Insignificant (i.e., not significantly different support) values are marked by ‘X’s. ‘No Modules’ refers to the null hypothesis of complete integration.

